# SciCore-Omics: a tri-modal foundation model unifying histology, spatial transcriptomics and language for spatial biology

**DOI:** 10.64898/2026.05.30.728937

**Authors:** Xinyu Xiao, Yunfei Li, Zheni Zeng, Yukun Yan, Zhiyuan Liu, Zhenghao Liu, Yujia Xiang, Zhenbang Ye, Jianming Ying, Yang Li, Linhai Xie, Fuchu He

## Abstract

Histomorphology and spatial transcriptomics capture complementary aspects of tissue biology, but their relationships remain difficult to extract, align, and interpret at scale. Existing foundation models typically connect histology, omics, or language only pairwise, which limits their capacity to jointly infer molecular states, decode spatial tissue organization, and generate biologically grounded explanations. Here, we show **SciCore-Omics**, the first tri-modal foundation model linking histology images, spatial transcriptomics, and biological language. We constructed a spatially paired image–gene–text dataset comprising 151,182 spots across multiple tissues and performed a three-stage progressive training of SciCore-Omics on this dataset. Across gene expression prediction and spatial domain recognition, SciCore-Omics achieved 23.6–80.9% relative gains in task-specific metrics over the strongest external baselines. It further showed robust zero-shot generalization in histopathology classification, outperforming GPT-5 by 6.16 percentage points in mean accuracy across four benchmarks. Expert evaluation in 10 breast cancer cases confirmed its H&E-only case-level molecular reasoning capability. Together, our method demonstrates that a tri-modal framework can effectively bridge histomorphology and molecular state, providing a more general and interpretable foundation model for computational pathology and omics analysis.

---

Histological images, particularly haematoxylin and eosin (H&E)-stained sections, provide rich information on tissue architecture, cellular organization and pathological alterations [1–3]. Spatial transcriptomics technologies capture local gene expression while preserving spatial context, offering a molecular view of tissue organization [4–7]. These two modalities provide complementary perspectives on the same biological system: histology reveals tissue morphology, while spatial transcriptomics reveals molecular programmes that underlie those phenotypes. Integrating morphological and molecular information is therefore a fundamental challenge in spatial biology and is essential for understanding tissue development, disease progression, therapeutic response and disease diagnosis.

Foundation models are rapidly advancing computational pathology and omics. Large-scale models trained on histopathology images [8, 9], transcriptomic data [10–12], or multimodal biomedical resources [13–25] have demonstrated remarkable abilities in representation learning and downstream prediction. However, most current approaches remain fragmented across modalities and are primarily optimized for predictive performance rather than biological reasoning. Pathology vision–language models can describe tissue morphology but lack direct access to molecular states [13–17]. Transcriptome–language models can interpret gene-expression patterns but cannot connect them to tissue architecture [18, 19]. Image– omics models learn shared embeddings or predictive mappings, yet typically provide limited semantic explanations of how morphological features relate to underlying molecular programmes [20–25]. As a result, there remains no unified framework capable of jointly integrating morphology, molecular state and biological semantics into an interpretable reasoning process. Biological language offers a natural bridge between these modalities, providing a semantic interface through which tissue structures, gene functions, pathway activities and disease-relevant mechanisms can be connected and interpreted.

To address this, we introduce **SciCore-Omics**, a language-centred tri-modal biomedical foundation model that integrates histology images, spatial transcriptomics and biological language within a decoder-based generative architecture. We developed **molecularly grounded semantic pairing** to construct an image–gene–text training resource comprising 151,182 spatial spots across multiple tissues. Spatially aligned H&E image patches and spot-level gene expression profiles were linked to transcriptomics-derived biological descriptions generated from highly expressed genes, enriched pathways and tissue context. Building on this dataset, SciCore-Omics employs a **progressive three-stage training strategy** to sequentially learn histopathological image semantics, gene expression semantics and multimodal joint representations, thereby achieving effective alignment across different modalities.

We systematically evaluated the performance of SciCore-Omics across various biomedical tasks. In gene expression prediction, SciCore-Omics increased the Pearson correlation coefficient (PCC) by 23.6% relative to OmiCLIP and reduced mean squared error (MSE) by 15.8%. For spatial domain detection, it achieved a spot-level macro-F1 of 0.850, representing an 80.9% relative gain over the strongest external embedding baseline, while also providing a joint image–gene generative interface. To assess whether tri-modal alignment improves generalizable tissue-level recognition without task-specific fine-tuning, we evaluated zero-shot histopathology classification across four H&E benchmarks, on which SciCore-Omics outperformed state-of-the-art models in mean accuracy. Furthermore, in a proof-of-concept clinical case-level reasoning evaluation involving 10 breast cancer cases, it attained the highest expert-rated reasoning score.

In summary, these results demonstrate the feasibility of unifying histomorphology, molecular state and biological language through tri-modal generative modelling, and establish SciCore-Omics as a general framework for integrative computational pathology and spatial omics analysis, with potential to support molecularly informed diagnosis and pathology decision-making.

## 1 Results

### 1.1 Overview of SciCore-Omics

We developed SciCore-Omics, a language-centred multimodal foundation model that unifies histology, spatial gene expression, and natural language within a shared token space (Fig. 1a,b). Specifically, the input histological images and gene expression profiles are processed by modality-specific encoders. Visual features are converted into a fixed number of vision virtual tokens through a visual encoder and a resampler module. Gene expression profiles are transformed into gene virtual tokens via a gene encoder, a Q-Former module, and a projection layer. The virtual tokens are then projected into the hidden dimension of the language model and inserted into placeholders, thereby forming a unified input embedding sequence. This sequence is further fed into the generative language backbone for autoregressive decoding. This design allows SciCore-Omics to treat histomorphology, molecular state, and user instruction as complementary contextual information in the same generative backbone, rather than as separate task-specific pipelines.

**Figure 1:**
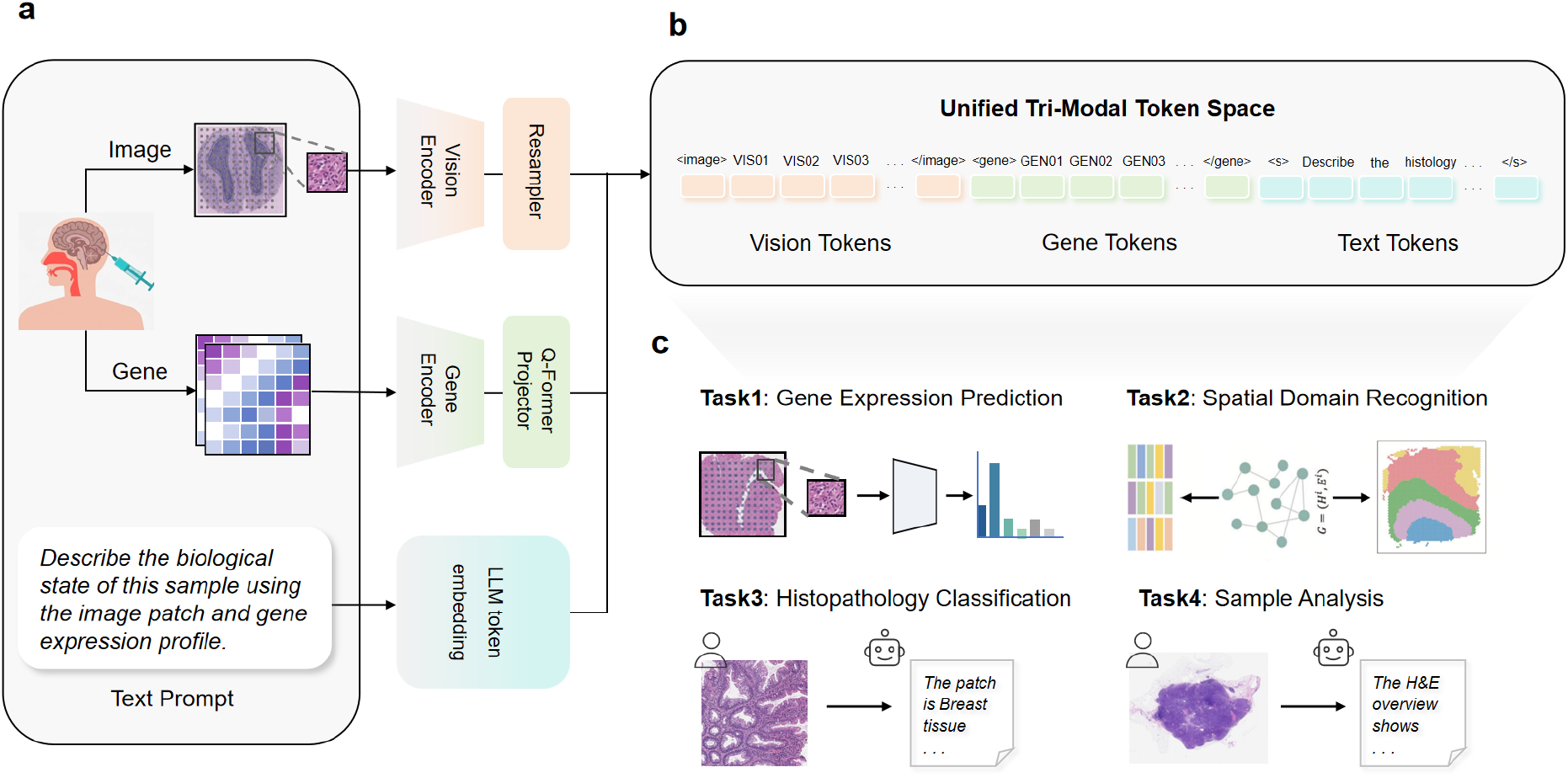
Overview of SciCore-Omics. **a**, Model architecture of SciCore-Omics. Histological images and spatial gene expression profiles are encoded by modality-specific encoders, and projected into the language-model space. **b**, Unified tri-modal token representation. Image, gene virtual tokens and text tokens are jointly modelled in a shared decoder-based backbone. **c**, Representative downstream tasks supported by SciCore-Omics across multiple biological scales.

The key advantage of tri-modal fusion at the token level is its ability to flexibly adapt to various forms of conditional input and support a wide range of multi-scale downstream tasks. Within the same underlying framework, SciCore-Omics can be applied to molecular-level gene expression prediction, spot-level spatial domain recognition, tissue-level pathology classification and sample-level analysis (Fig. 1c).

### 1.2 Progressive multimodal alignment improves transcriptome-to-language

Spatial transcriptomics technology integrates histological imaging with high-throughput sequencing to provide gene expression profiles corresponding to histological sections. However, the interpretation of these data relies heavily on annotation, and manual annotation is slow, subjective, and difficult to scale to large numbers of spots. Therefore, we deployed the molecularly grounded semantic pairing pipeline to automatically extract structured biological evidence directly from the gene expression profiles and tissue metadata of each spot, including tissue origin, highly expressed marker genes, and enriched pathways obtained from single-sample gene set enrichment analysis (ssGSEA) [26–29]. We used this structured information to generate richer biologically grounded descriptions (Fig. 2a), thereby constructing image– gene–text triplets. This enables the model not only to learn the correspondence between histological images and genes but also to articulate complex biological characteristics using language.

**Figure 2:**
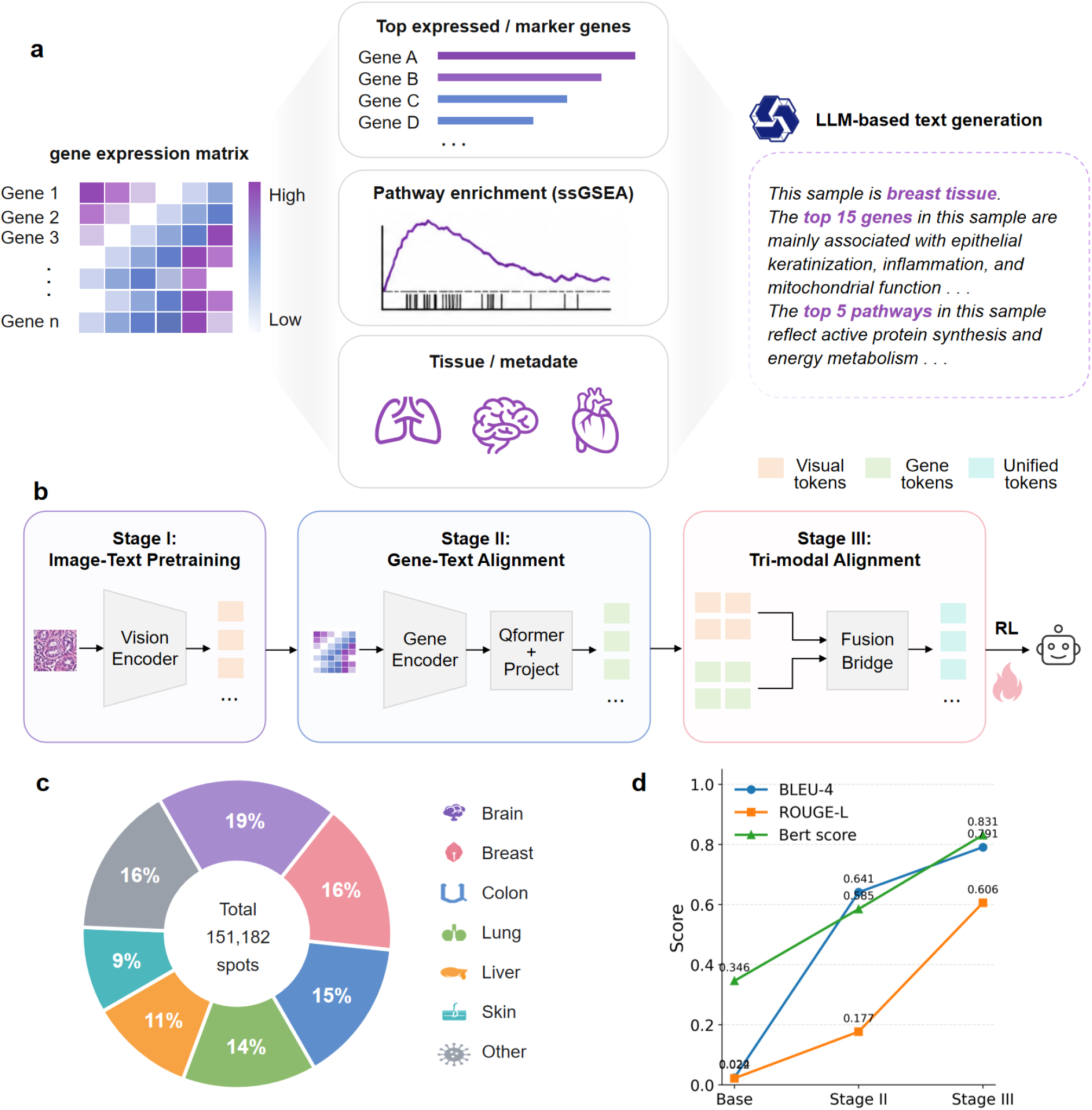
Progressive multimodal alignment of SciCore-Omics. **a**, Transcriptome-grounded descriptions are generated from spot-level gene expression, marker genes, enriched pathways and tissue metadata. **b**, Three-stage training strategy comprising image–text pretraining, gene–text alignment and tri-modal alignment, followed by lightweight reinforcement learning. **c**, Summary of the pretraining corpus and tissue distribution of the curated spatial transcriptomics data. **d**, Transcriptome-to-language generation performance of models from different training stages on a held-out validation split of the gene–text paired dataset, evaluated using BLEU-4, ROUGE-L and BERTScore F1.

We adopted an incremental training strategy to link histological images, transcriptomic data, and linguistic information in phases (Fig. 2b and Extended Data Fig. 1). In the first stage, we pre-trained a basic vision–language foundation model to learn pathology-related visual features and semantic information. In the second stage, we incorporated a gene encoder and a projection module to align transcriptomic representations with the language latent space. In the third stage, we performed joint optimization of the model’s representations using tri-modal paired data. Additionally, we applied lightweight reinforcement learning to ensure the standardization of the model’s output format and the consistency of its linguistic expressions. By collecting public datasets and data from existing studies, we constructed a large-scale pre-training corpus for the three-stage training, comprising 1,052,334 images, approximately 39.85 million text sentences, and 151,182 tri-modal data pairs, covering various tissue types such as brain, breast, colon, lung, liver, and skin (Fig. 2c and Extended Data Fig. 2).

To validate whether progressive training improved transcriptome-conditioned language generation, we evaluated models from different training stages on a held-out validation split of the gene–text paired dataset that was not used for model training (Fig. 2d). Given transcriptomic input, each model was prompted to generate a biological description, which was compared with the corresponding transcriptome-derived reference description. Text generation quality was evaluated using complementary metrics. BLEU-4 and ROUGE-L measure local lexical overlap and sequence structural similarity between generated text and reference descriptions, respectively [30, 31]. Biomedical semantic fidelity was assessed using BERTScore F1, which measures embedding-level semantic similarity between generated and reference descriptions in a biomedical language space [32]. Compared to the baseline model without explicit gene–text alignment, the second stage improved BLEU-4 from 0.024 to 0.641, ROUGE-L from 0.022 to 0.177, and BERTScore from 0.346 to 0.585. The performance of the third-stage model improved further, achieving a BLEU-4 score of 0.791, a ROUGE-L score of 0.606, and a BERTScore of 0.831. These results indicate that phased multimodal alignment significantly enhances the model’s ability to generate descriptions from transcriptomic data, with the generated descriptions exhibiting high lexical overlap, strong structural consistency, and excellent biomedical semantic fidelity.

### 1.3 SciCore-Omics predicts gene expression from histology

Spatial expression prediction is a critical task in computational biology, aimed at inferring gene expression patterns without relying on expensive spatial transcriptomics sequencing. SciCore-Omics predicts gene expression by extracting features from H&E images via a visual encoder (Fig. 3a). The use of AI models to infer gene expression from H&E images has been explored in several recent studies [21–25]. We conducted a systematic comparison of SciCore-Omics with existing methods on a dataset of normal human heart samples [33]. MSE and PCC were used as performance metrics.

**Figure 3:**
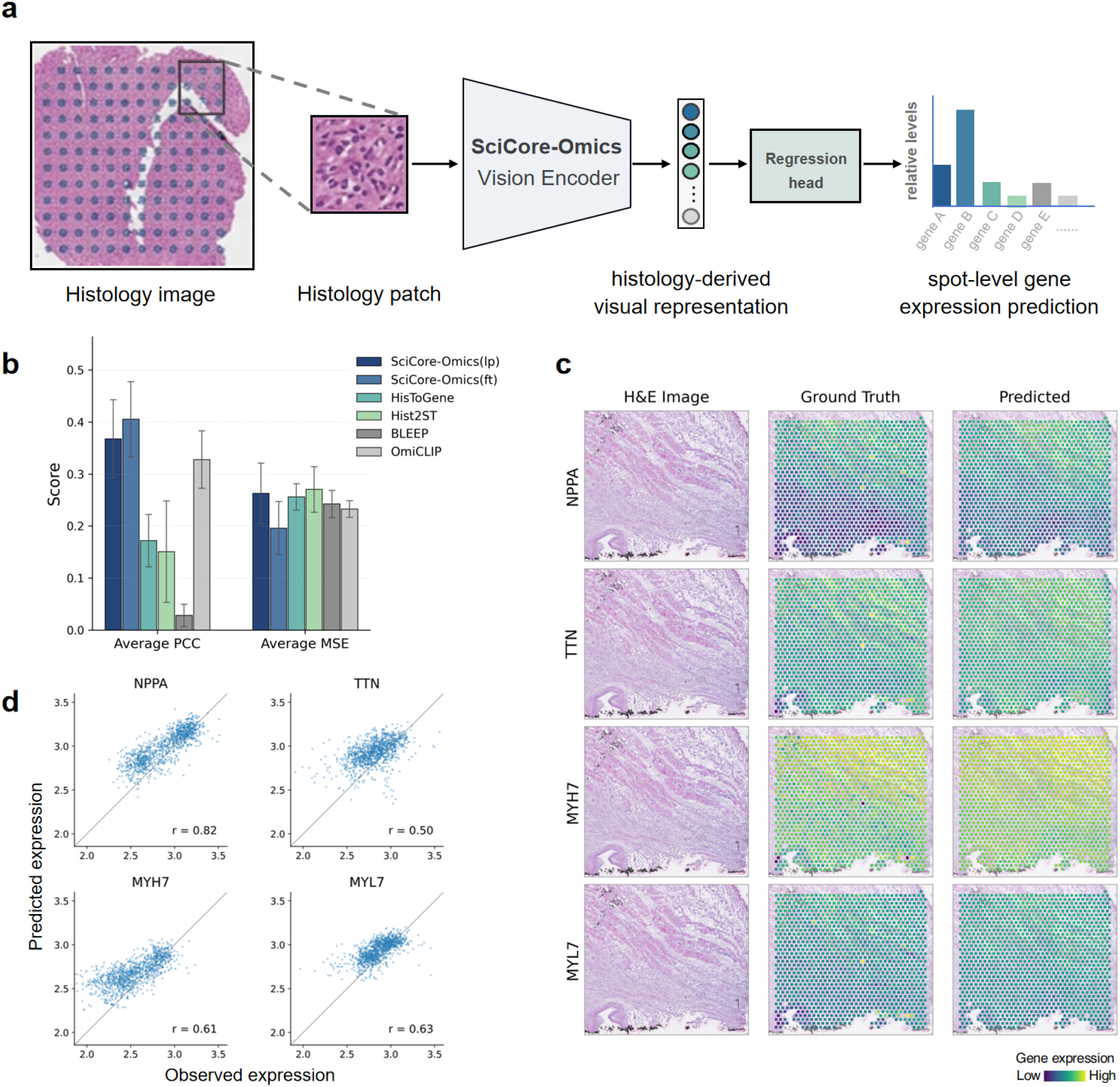
Histology-based gene expression prediction with SciCore-Omics. **a**, Schematic illustration of molecular-level gene expression prediction. H&E image patches are encoded by the SciCore-Omics vision encoder, followed by a regression head to predict spot-level gene expression profiles. **b**, Performance comparison on 39 normal human heart spatial transcriptomics sections using PCC and MSE. LP, linear probing with a frozen visual encoder; FT, fine-tuning. Error bars indicate s.d. **c**, Representative measured and predicted spatial expression maps. **d**, Scatterplots comparing observed and predicted expression values for representative genes, with Pearson correlations shown.

Compared to OmiCLIP, Hist2ST, HisToGene, and BLEEP, SciCore-Omics achieved the best MSE performance and the highest mean PCC (Fig. 3b and Extended Data Fig. 3). Under the frozen visual encoder setting, the model achieved an average PCC of 0.3679 and an average MSE of 0.2629. In the fine-tuned setting, SciCore-Omics further improved performance, increasing mean PCC by 23.6% and reducing mean MSE by 15.8% relative to OmiCLIP. These results indicate that the visual representations learned by SciCore-Omics contain rich molecular information and can be further adapted to downstream prediction tasks through task-specific fine-tuning.

This was further illustrated by the spatial distribution prediction results for representative genes (Fig. 3c). For the gene *NPPA*, which exhibits distinct spatial confinement in cardiac tissue, the predicted expression map aligns highly with the observed distribution, with a correlation of 0.82 at the spot level. For *TTN* and *MYH7*, SciCore-Omics successfully predicted their correspondence with tissue regional structures, with correlations of 0.50 and 0.61, respectively. Similar spatial consistency was also observed for *MYL7* (*r* = 0.63). In these examples, the predicted maps effectively reproduce the major spatial features observed in the real expression data. However, compared to the measured expression values, the predicted results exhibit a certain degree of compression, resulting in a smoother overall profile, consistent with the scatterplot results shown in Fig. 3d.

Together, SciCore-Omics is robust in predicting gene expression. Compared to methods such as OmiCLIP, which rely on contrastive learning frameworks and retrieve transcriptomic profiles from a reference set, SciCore-Omics, as a generative model, can directly generate transcriptomic profiles, offering greater accessibility and scalability.

### 1.4 SciCore-Omics improves spatial domain recognition

Another important task in spatial transcriptomics analysis is spatial domain identification [34–36]. We assessed SciCore-Omics in two settings. First, we examined whether the embeddings learned by the model encode spatial domain information by training a classification head on a frozen model to predict spatial domain labels. Second, we assessed whether the model can generate natural language descriptions containing spatial domain-related information. We defined the task as follows: for each spot, SciCore-Omics takes histological image patches and/or gene expression profiles as input to generate biological descriptions from which region labels, pathway labels, and marker genes are extracted (Fig. 4a). We conducted evaluations on a spatial transcriptomics dataset of the human dorsolateral prefrontal cortex (DLPFC) [37]. This dataset comprises 12 tissue sections with paired H&E images and spatial gene expression data generated by the 10x Genomics Visium platform, along with manual cortical layer annotations.

**Figure 4:**
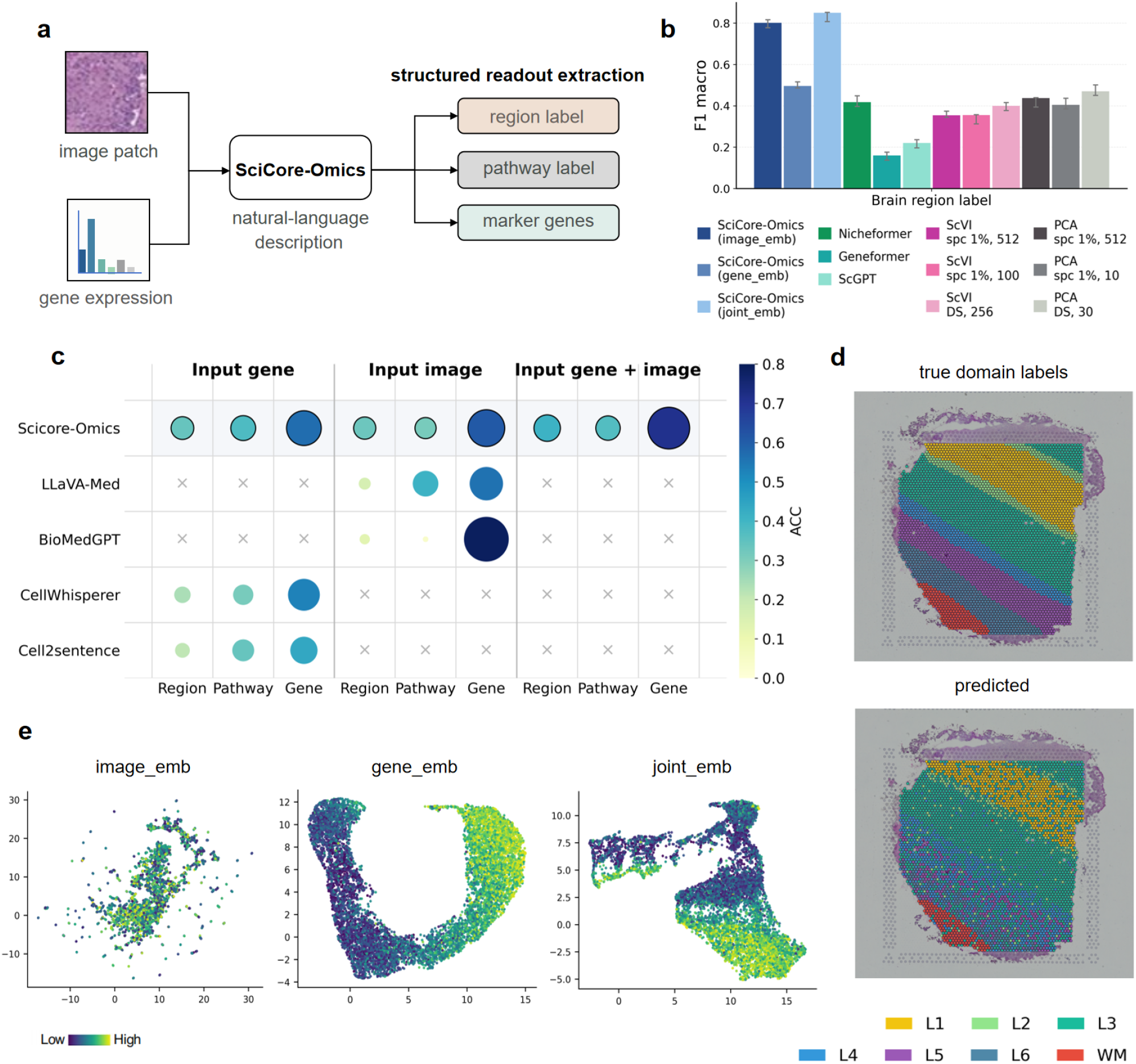
Spot-level spatial domain recognition in the human DLPFC. **a**, Schematic illustration of spot-level spatial domain recognition. For each spot, SciCore-Omics takes the matched histology image patch and/or gene expression profile as input and generates a description, from which region labels, pathway labels and marker genes are extracted. **b**, Test-set macro-F1 scores for brain-region label prediction using linear-probing models computed from frozen embeddings generated by SciCore-Omics, Nicheformer, Geneformer, scGPT, scVI and PCA. **c**, Accuracy of region, pathway and marker-gene prediction for models after fine-tuning. Bubble size and colour indicate ACC for each structured readout. Grey crosses indicate unsupported input–output settings. **d**, Spatial projection of manual and predicted brain-region labels in held-out DLPFC sections. **e**, UMAP visualization of image-only, gene-only and joint image–gene embeddings coloured by *Glutamatergic_synapse* pathway activity.

In the first experiment, SciCore-Omics achieved the highest macro-F1 score (Fig. 4b). Notably, compared with image-only embeddings (0.801) and gene-only embeddings (0.496), joint image–gene embeddings further improved the macro-F1 score to 0.850 [10–12, 38]. This corresponded to an 80.9% relative improvement over the strongest external embedding baseline among Nicheformer, Geneformer, scGPT, scVI and PCA (macro-F1 = 0.470), highlighting the added value of multimodal representation learning for spatial tissue organization.

In the second experiment, we selected eight tissue sections to construct a small-scale tri-modal paired dataset for model fine-tuning and evaluated the model on the remaining four held-out sections. Accuracy (ACC) was used as the evaluation metric, and the comparison with representative biomedical multimodal foundation models is shown in Fig. 4c. Under gene-only input, SciCore-Omics achieved a mean ACC of 0.430 across region, pathway and gene prediction, outperforming the gene-input baselines CellWhisperer and Cell2Sentence by 19.4% and 30.3%, respectively. Under image-only input, SciCore-Omics achieved a mean ACC of 0.417, exceeding LLaVA-Med and BioMedGPT by 12.6% and 30.2%, respectively. Although some baselines performed better on individual subtasks, their performance was less balanced across output types. When image and gene inputs were jointly provided, SciCore-Omics achieved the highest mean ACC of 0.497, with ACC values of 0.41, 0.37 and 0.71 for region, pathway and gene prediction, respectively. Joint image–gene conditioning improved mean ACC by 15.5% over the gene-only setting and by 19.2% over the image-only setting, indicating that integrating molecular and morphological information improves balanced spatial biological prediction.

To assess whether the prediction results accurately reflect spatial structures, we remapped the predicted regional labels back onto the original tissue sections. The predicted map generally recovered the typical cortical layering structure in the DLPFC and preserved the large-scale spatial arrangement of major regions within the section (Fig. 4d). Prediction errors primarily occurred near the boundaries of adjacent regions, where there are typically continuous morphological transitions and a mixing of transcriptional states between neighbouring spots. We further projected the embeddings into UMAP space and coloured spots according to pathway activity scores (Fig. 4e and Extended Data Fig. 4). These visualizations revealed pathway-specific organization in the learned embedding spaces: spots with high pathway activity tended to occupy locally enriched regions or continuous gradients rather than being randomly distributed. In the representative *Glutamatergic synapse* pathway shown in Fig. 4e, the gene-derived embedding showed the strongest organization of high-activity spots, consistent with the transcriptional nature of this synaptic program, while the joint image–gene embedding retained a comparable but less pronounced pathway-associated structure. In contrast, morphology-associated pathways such as *Myelination* and *Oligodendrocyte differentiation* were more clearly organized in the image and joint embedding spaces. Similar modality-dependent patterns were observed for additional brain-related pathways, including *Axonogenesis* and *Neuron projection* (Extended Data Fig. 4).

### 1.5 SciCore-Omics supports zero-shot histopathology classification

Next, we evaluated SciCore-Omics’ performance in zero-shot histopathological classification. We assessed the model in a zero-shot, closed-set classification setting across four public H&E benchmarks: MHIST, BACH, CRC100K and PatchCamelyon, which cover distinct histopathology classification scenarios [39– 43]. For each image, the prompt specified the dataset-specific candidate labels and instructed the model to select one category without task-specific fine-tuning or in-context examples (Fig. 5a).

**Figure 5:**
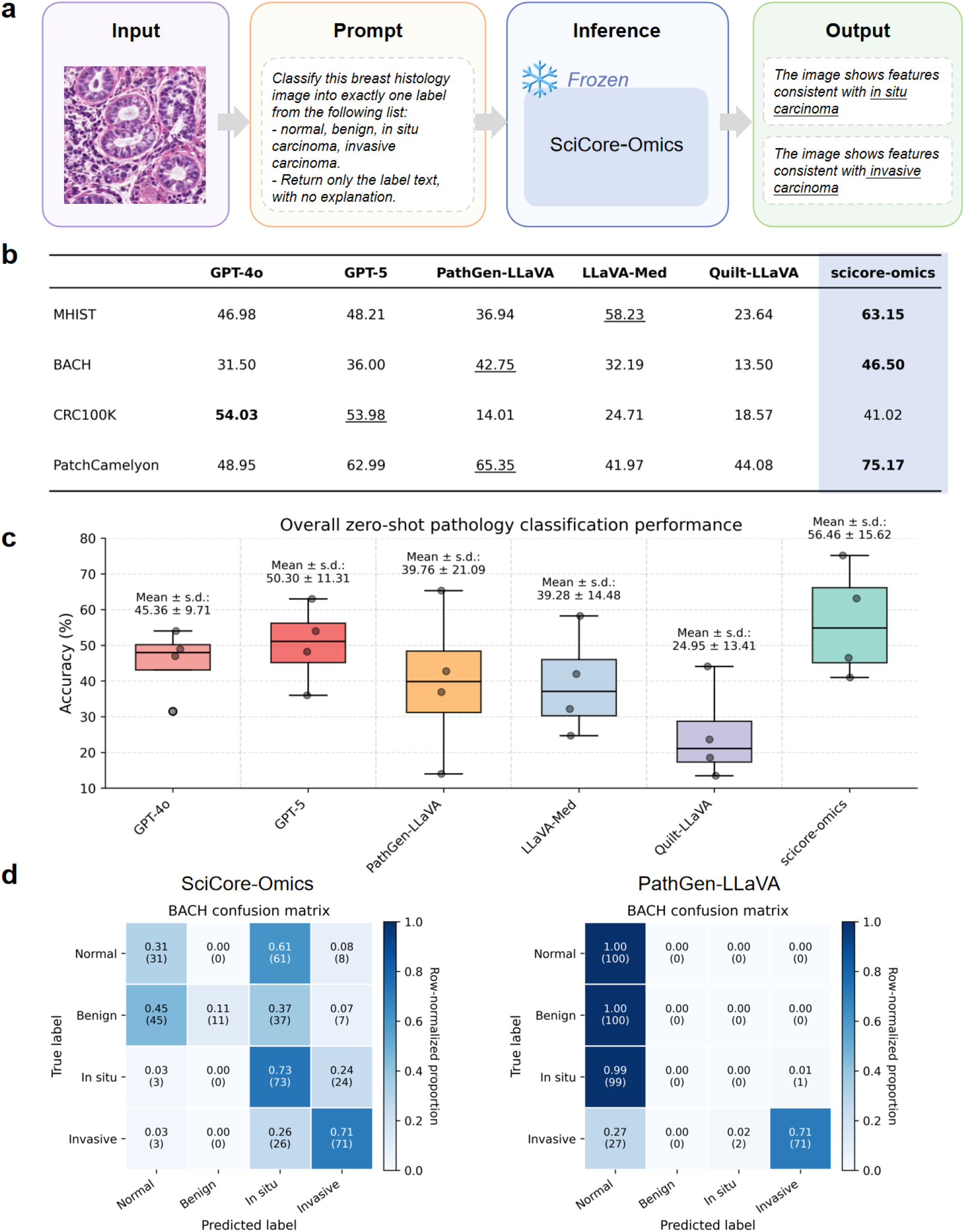
Zero-shot histopathology classification. **a**, Closed-set zero-shot evaluation using dataset-specific class prompts. **b**, Accuracy on MHIST, BACH, CRC100K and PatchCamelyon compared with general and biomedical VLM baselines; bold and underlining indicate best and second-best results. **c**, Accuracy distribution across the four benchmarks for each model. Centre values indicate mean accuracy, and error bars indicate s.d. across benchmarks (n = 4). **d**, Row-normalized BACH confusion matrices for SciCore-Omics and PathGen-LLaVA, with raw counts in parentheses.

SciCore-Omics achieved the highest accuracy on three of the four benchmarks, reaching 63.15% on MHIST, 46.50% on BACH and 75.17% on PatchCamelyon (Fig. 5b). On CRC100K, GPT-4o and GPT-5 achieved higher accuracies of 54.03% and 53.98%, while SciCore-Omics obtained 41.02%. Nevertheless, SciCore-Omics remained above the evaluated biomedical vision–language model (VLM) baselines on this dataset, including PathGen-LLaVA, LLaVA-Med and Quilt-LLaVA [15–17]. Across repeated evaluations on the four benchmarks, SciCore-Omics achieved the highest macro-averaged accuracy, with a mean accuracy of 56.46% across benchmarks, compared with 50.30% for GPT-5, 45.36% for GPT-4o, 39.76% for PathGen-LLaVA, 39.28% for LLaVA-Med and 24.95% for Quilt-LLaVA (Fig. 5c). BACH confusion matrices showed that SciCore-Omics produced more balanced predictions than PathGen-LLaVA, which was strongly biased towards the normal class (Fig. 5d). SciCore-Omics improved recall for in situ and invasive carcinoma, although confusion among normal, benign and in situ carcinoma cases persisted. Together, SciCore-Omics retains transferable tissue-level visual recognition ability within a broader visual–omics–language framework.

### 1.6 SciCore-Omics enables H&E-based pathology reasoning

To further evaluate the pathology reasoning capability and potential clinical applicability of SciCore-Omics, we assessed the model in both patch-level and case-level H&E-based settings. We first evaluated SciCore-Omics on PathVQA, a pathology visual question answering benchmark consisting of histopathology images paired with medically grounded questions and answers [44]. In the fine-tuned setting, SciCore-Omics achieved an open-ended score of 38.10 and a closed-ended score of 88.66 (Fig. 6a). Compared with representative MedVQA and biomedical VLM baselines, including MMQ, M2I2 and BiomedGPT, SciCore-Omics achieved competitive performance on closed-ended questions and obtained the highest open-ended score among the compared models.

**Figure 6:**
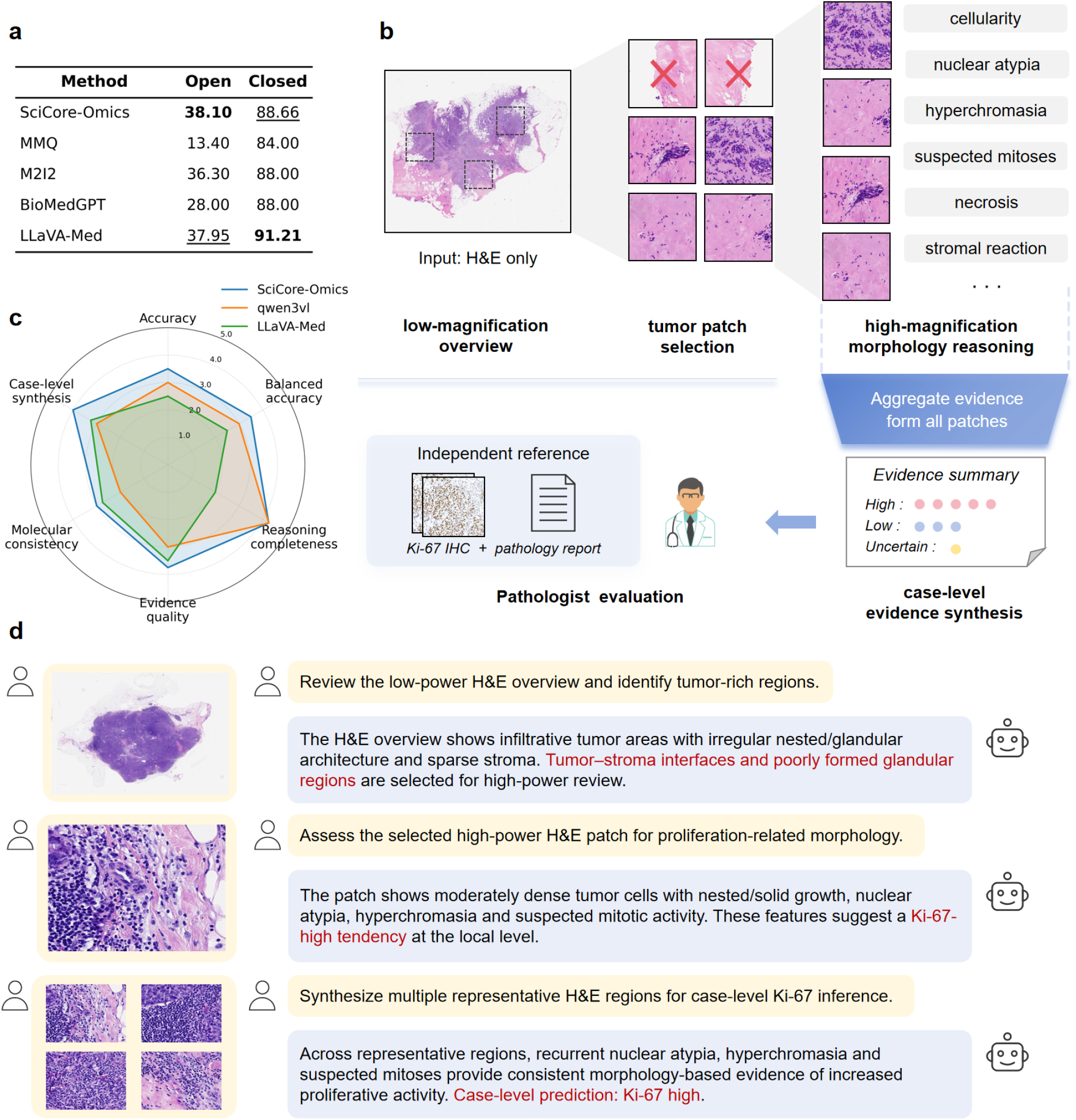
H&E-based pathology reasoning with SciCore-Omics. **a**, Fine-tuned PathVQA performance on open-ended and closed-ended questions compared with biomedical VLM baselines. **b**, H&E-only case-level reasoning workflow, from low-power slide review to high-power patch assessment and case-level synthesis; Ki-67 immunohistochemistry and pathology reports were used only for expert evaluation. **c**, Expert assessment of SciCore-Omics, Qwen3-VL and LLaVA-Med on 10 breast cancer cases across six reasoning dimensions. **d**, Representative proliferation-state inference from tumour-rich regions, high-power morphology and aggregated case-level evidence.

We next examined whether the model could extend this capability to a more clinically oriented case-level reasoning setting. In this evaluation, only the whole-slide H&E images were used as the model input, while immunohistochemistry and pathology reports were withheld from the model and referenced solely for expert assessment (Fig. 6b). The inference workflow was designed to simulate a staged pathological review process. First, SciCore-Omics reviewed the low-magnification overview of the whole-slide image to identify tumour-rich regions and representative tumour–stroma structures. Second, representative tumour patches were selected from these regions for high-magnification assessment, with attention to interpretable morphological features, including tumour cell density, nuclear atypia, hyperchromasia, suspected mitotic activity, necrosis and stromal reaction. Finally, the model integrated evidence across multiple regions to generate a case-level interpretation of the proliferative status.

Clinical experts assessed the outputs of SciCore-Omics, Qwen3-VL and LLaVA-Med on 10 breast cancer cases. The evaluation covered accuracy, balanced accuracy, reasoning completeness, evidence quality, molecular consistency and case-level synthesis (Fig. 6c), thus emphasizing both final clinical consistency and the plausibility of evidence and synthesis. SciCore-Omics achieved the highest overall mean expert score among the compared models, with an average score of 3.67 compared with 3.04 for Qwen3-VL and 2.75 for LLaVA-Med. Across individual dimensions, SciCore-Omics achieved the highest scores in accuracy, balanced accuracy, evidence quality, molecular consistency and case-level synthesis, and tied with Qwen3-VL in reasoning completeness.

A representative case further illustrates the staged reasoning process of SciCore-Omics (Fig. 6d and Extended Data Fig. 5). From the low-power H&E overview, the model identified infiltrative tumour-rich regions and prioritized tumour–stroma interfaces and poorly formed glandular areas for high-power review. In the selected high-power regions, it described moderately dense tumour cells with nested or solid growth, nuclear atypia, hyperchromasia and suspected mitotic activity. By integrating recurrent findings across representative regions, SciCore-Omics generated a case-level interpretation of a Ki-67-high proliferative state, which was consistent with the independent immunohistochemistry and pathology-report references.

Because the expert evaluation was performed on a limited number of cases, these results should be interpreted as a clinically oriented case study rather than a definitive clinical validation. Nevertheless, the assessment provides initial evidence that SciCore-Omics can support H&E-based pathology reasoning by integrating morphological findings across tumour regions and generating interpretable case-level inferences consistent with independent molecular references.

## 2 Discussion

Existing studies have largely advanced along separate trajectories, including histology–omics, histology– language and omics–language modelling [15, 18, 19, 22–24]. This leaves a critical gap: a unified generative framework in which molecular and morphological evidence jointly guides language generation. SciCore-Omics fills this gap by projecting histomorphology, spatial transcriptomics and biomedical language into a shared autoregressive token space, thereby establishing the first tri-modal foundation model centred on language generation.

Our results support three main conclusions. First and most importantly, transcriptome-to-language supervision provides a scalable, annotation-free strategy for omics–language alignment, demonstrating that structured evidence from spatial transcriptomics can be directly translated into natural language. Second, the tri-modal architecture is inherently multi-task: within a unified framework, it supports transcriptome-conditioned generation, molecular prediction, spatial domain recognition, tissue classification, pathology question answering, and case-level assessment. Third, the model offers an interpretable semantic workspace where image-derived structures and transcriptome-derived signals jointly inform biologically grounded hypotheses.

Several limitations should be noted alongside these strengths. First, our evaluation spans diverse biological contexts rather than being structured around a single disease-centred axis. While this supports assessment of cross-modal generalizability, it limits our ability to establish a closed-loop biological narrative from molecular state to tissue phenotype within one disease type. Second, the training corpus relies primarily on tissue-category labels and transcriptome-derived descriptions, with limited fine-grained annotations (e.g., cell-type composition, pathological subtypes, clinically actionable labels). Third, the ten-case pathology workflow remains a proof-of-concept; the retrospective design and small sample size preclude definitive conclusions about clinical utility.

Future work will focus on three directions. First, we will perform closed-loop tri-modal validation in a well-defined single-disease cohort (e.g., breast or lung cancer), jointly evaluating histomorphology, spatial transcriptomics, immunohistochemistry and pathology reports. Second, we will construct finer-grained label corpora to enable cell-type annotation, pathological subtyping and clinically actionable molecular reasoning. Third, we will scale the pathology workflow to prospective whole-slide studies with hundreds of cases, incorporating expert review and clinically grounded evaluation protocols.

## 3 Online Methods

### 3.1 Study design and evaluation overview

This study was designed to develop and evaluate SciCore-Omics, a tri-modal foundation model that connects histology images, spatially resolved gene expression profiles and biological language within a unified generative framework. The primary unit for multimodal alignment was the spatial transcriptomics spot. When paired histology and spatial transcriptomics data were available, each spot was represented by a local H&E image patch, a matched gene expression vector and, for language-supervised training, a transcriptome-grounded biological description derived from structured molecular evidence.

Model performance was assessed across complementary levels of biological abstraction. Molecular-level evaluation tested histology-based spatial gene expression prediction. Spot-level evaluation tested spatial domain recognition, pathway recovery and marker-gene recovery. Tissue-level evaluation tested zero-shot histopathology classification on public H&E benchmarks. Language-based evaluation tested transcriptome-conditioned biological description generation and pathology visual question answering. Finally, a pilot whole-slide workflow evaluated whether H&E-only case-level reasoning could produce interpretable proliferation-related assessments in breast cancer cases.

For transcriptome-to-language generation, we used held-out spatial transcriptomics samples with paired gene expression profiles and reference descriptions. For molecular-level spatial gene expression prediction, we used the Human Heart Cell Atlas spatial transcriptomics dataset, which comprised 39 normal human heart spatial transcriptomics sections [33]. For spot-level spatial domain recognition, we used the human dorsolateral prefrontal cortex spatial transcriptomics dataset, which comprised 12 sections with paired H&E images, gene expression profiles and manual cortical-layer annotations [37]. For zero-shot tissue-level histopathology classification, we used MHIST, BACH, NCT-CRC-HE-100K with CRC-VAL-HE-7K and PatchCamelyon [39–43]. For pathology visual question answering, we used PathVQA [44]. For the pilot whole-slide pathology workflow, we used an internal cohort of 10 breast cancer cases with H&E whole-slide images, Ki-67 immunohistochemistry and pathology-report references. The immunohistochemistry and reports were used only for expert evaluation and were not provided to the model as input.

### 3.2 Ethics approval

The use of the internal breast cancer cohort, including H&E whole-slide images, Ki-67 immunohistochemistry images and pathology-report references, was reviewed and approved by the Ethics Committee of National Cancer Center/Cancer Hospital, Chinese Academy of Medical Sciences and Peking Union Medical College (approval no. 25/077-5023). All procedures involving human-derived data were conducted in accordance with the relevant institutional guidelines and regulations. The cohort was retrospectively analysed using de-identified data. The requirement for written informed consent was waived by the ethics committee owing to the retrospective and de-identified nature of the study.

### 3.3 Model architecture

SciCore-Omics is a language-centred multimodal foundation model built on MiniCPM-V as the generative multimodal language backbone and Nicheformer as the gene encoder [12, 45–47]. Histology images and gene expression profiles are processed by modality-specific encoders and then projected into the hidden dimension of the language model. The resulting visual and molecular token spans replace reserved modality placeholders in the input sequence, allowing image, gene and text tokens to be jointly processed by the same decoder-based generative backbone.

The complete SciCore-Omics model used in this study contains approximately **8.5 billion parameters**, including the language-centred multimodal backbone, the histology visual branch, the Nicheformer-based gene encoder, the Q-Former module and modality projection layers. This parameter scale provides sufficient capacity for tri-modal generative reasoning over histomorphology, spatial transcriptomics and biological language, while remaining compatible with controlled local deployment on institutional GPU servers.

In the implementation used in this study, the gene branch follows

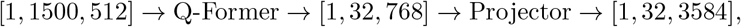

where the gene encoder first maps the gene expression profile into molecular embeddings, the Q-Former compresses the molecular representation into 32 query virtual tokens, and the projector maps these virtual tokens into the language-model hidden space. The visual branch follows

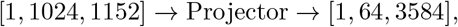

where histology-derived visual features are compressed into 64 visual virtual tokens and projected into the same token space as the molecular virtual tokens. The final multimodal sequence consists of text tokens, optional visual virtual tokens and optional gene virtual tokens, followed by answer tokens for autoregressive decoding.

This design allows SciCore-Omics to operate under image-only, gene-only or joint image–gene input settings without requiring task-specific architectural changes. For generative tasks, the language model decodes free-form or structured biological text conditioned on the available modalities. For discriminative tasks, the same model can be prompted in a closed-set manner or used as a feature extractor for lightweight downstream heads.

### 3.4 Spot-level preprocessing and transcriptome-to-language supervision

For spatial transcriptomics samples, spot coordinates were mapped to the paired H&E image using the coordinate metadata supplied with each dataset. A square histology patch centred on each spot was extracted from the corresponding tissue image and resized to the input resolution required by the visual encoder. Patch extraction followed coordinate registration so that each local image patch represented the same spatial region as its matched gene expression profile. When tissue masks were provided by the dataset, patches with substantial background or tissue-fold artefacts were excluded using the available masks. When tissue masks were unavailable, background filtering was performed using intensity- and saturation-based heuristics followed by manual spot checks on representative slides. Image preprocessing was performed within each dataset before model input and did not use information from held-out evaluation labels. Dataset-specific patch extraction settings, including image scale, patch size and input resolution, are provided in the released preprocessing configuration.

Gene expression matrices were processed separately for each dataset before being used as model input. For count-based spatial transcriptomics data, raw counts were normalized for library size, log-transformed and mapped to a common gene-symbol vocabulary whenever possible. Genes outside the gene encoder vocabulary were omitted from model input. When benchmark-specific gene sets were required, gene selection was performed using the training split only, unless the benchmark provided a fixed predefined gene list. The processed expression vector was converted into the fixed-dimensional representation expected by the gene encoder. For downstream gene expression prediction, the target expression values were retained according to the benchmark protocol and evaluated only on held-out sections. For pathway and marker-gene evaluation, pathway activity and marker-gene labels were derived from structured expression-based analyses rather than from free-form model outputs.

To construct language supervision for spatial transcriptomics spots, each spot *i* was represented as a tuple

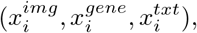

where 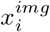 denotes the local histology patch, 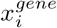 denotes the matched gene expression profile and 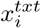 denotes a natural-language biological description derived from structured evidence associated with the same transcriptomic measurement.

Text supervision was generated from structured biological evidence rather than from unconstrained language-model speculation. For each spot, molecular evidence was summarized using highly expressed genes, representative marker genes, tissue metadata and pathway-enrichment labels. Pathway activity was computed using single-sample gene set enrichment analysis with curated pathway gene sets, including KEGG gene sets [26–29]. These structured fields were inserted into an evidence-constrained prompt and verbalized into concise biological descriptions using GPT-4 [48].

The language model was used only to verbalize precomputed structured evidence. It was instructed to preserve supplied gene symbols, pathway names and tissue labels; to use only the provided evidence fields; and to avoid unsupported disease, cell-type or mechanistic claims. Therefore, the generated descriptions were treated as natural-language renderings of structured annotations rather than as independent biological ground truth.

Generated descriptions were filtered for missing required evidence fields, malformed gene symbols, unsupported tissue names, formatting failures and obvious violations of the evidence constraints. Descriptions passing quality control were used for Stage II gene–text alignment and Stage III tri-modal joint training.

### 3.5 Training corpus construction and progressive multimodal alignment

We constructed a multimodal training corpus that integrated text-only, image–text, gene–text and image– gene–text data. These datasets were organized into a progressive three-stage curriculum designed to align histology, transcriptomics and biological language in a stable manner. To reduce catastrophic forgetting during training, the mixed data in the third stage also retained general text-only and general vision– language data, such as UltraChat and Flickr30k [49, 50]. In total, the corpus contained approximately 1.05 million images, 39.85 million text sentences and 151,182 tri-modal data pairs. Dataset composition and statistics are provided in Supplementary Table 1 and Extended Data Fig. 2. The module-level implementation and trainable components of this progressive curriculum are illustrated in Extended Data Fig. 1.

#### Stage I: histology–text alignment

The first stage adapted the visual branch and language backbone to pathology image–text semantics. We used pathology textbooks and pathology image–text pairs collected from publicly available biomedical literature, including PubMed Central and pathology-oriented image–text corpora, yielding approximately 503,016 images and 28.52 million text sentences [13, 16]. This stage was implemented as pretraining to establish correspondence between tissue morphology, cellular appearance and pathology terminology [13–17].

#### Stage II: gene–text alignment

The second stage initialized the omics-to-language pathway. Approximately 51,602 gene–text pairs were constructed from public transcriptomic resources, including GEO and 10x Genomics datasets [4, 37, 51]. Gene expression profiles were converted into structured biological evidence, including highly expressed genes, pathway-enrichment labels and tissue metadata, and were further transformed into biological descriptions using the evidence-constrained procedure. This stage updated the gene encoder and projection modules so that transcriptomic representations could be mapped into the semantic space used by the language model.

#### Stage III: tri-modal joint alignment

The third stage performed spot-level image–gene–language alignment using spatial transcriptomics data with paired H&E images and gene expression profiles. After quality control, 151,182 spatial spots were used as tri-modal image–gene–text triplets. Each local image patch and matched gene expression vector were paired with a transcriptome-grounded description derived from the same spot. During this stage, the model was trained on image-only, gene-only and joint image– gene input formats to support flexible downstream use. General-domain image–text and instruction-following samples were mixed at a low proportion to preserve general multimodal and language-following competence.

### 3.6 Training objectives and offline reward-weighted refinement

#### Training objectives

The main alignment objective combined autoregressive generation with multi-modal contrastive alignment:

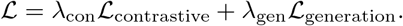

The generative term was the standard autoregressive language-modelling loss, optimizing the conditional likelihood of the target response given the available image, gene and text tokens:

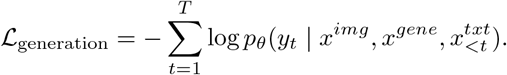

The contrastive term encouraged matched modalities from the same biological sample to be close while separating mismatched samples within the mini-batch [52, 53]. Pairwise InfoNCE-style losses were computed across available modality pairs, including image–text, gene–text and image–gene pairs when present [52, 53]. Modality-level representations were obtained from the corresponding projected token spans before contrastive comparison. The loss weights and temperature parameters were fixed within each training stage and are provided in the released training configuration.

#### Offline reward-weighted refinement

After the three-stage multimodal alignment curriculum, we applied a lightweight offline reward-weighted reinforcement learning step to improve answer formatting, language consistency and evidence grounding. This step was not designed as a core modality-alignment component and was not used to introduce new image–gene–language supervision. Instead, it was applied after tri-modal alignment to reduce unsupported biological claims, mixed Chinese–English outputs and failures to follow required output schemas.

We used an offline rather than online rollout setting. Candidate answers were not generated during reinforcement learning. Instead, each training instance contained a fixed response from curated instruction-style data, and the model was optimized using the token-level likelihood of this fixed response weighted by rule-based quality scores. This design was chosen to avoid instability from online response generation in a tri-modal setting and to focus reinforcement learning on response reliability rather than exploration.

For gene-conditioned examples, the molecular input was represented using the same 32-token gene span as in tri-modal training. When an instruction example contained a single gene placeholder, the placeholder was expanded to the molecular-token span produced by the gene projection module. The resulting sequence contained text tokens, optional image virtual tokens and optional gene virtual tokens, followed by the supervised answer span.

Rewards were rule-based rather than learned from a separately trained reward model. Each fixed response was assigned a scalar reward based on preprocessed quality scores and structured checks for format adherence, language consistency and evidence consistency. Responses were penalized if they omitted required fields, failed to follow the required schema, mixed languages inappropriately or introduced unsupported biological claims not grounded in the supplied genes, pathways, tissue labels or visual context.

For a prompt–response pair (*x, y*), where *x* denotes the multimodal input and *y* = (*y*_1_, … , *y*_*T*_) denotes the fixed answer span, token-level log probabilities were computed under the current policy *π*_*θ*_ and a reference policy *π*_old_. The reward *R*(*x, y*) was converted into an advantage term *A*(*x, y*), and the reinforcement-learning objective followed a clipped reward-weighted likelihood-ratio form:

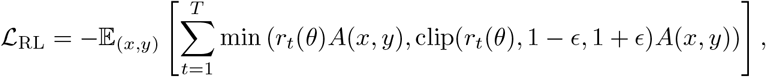

where

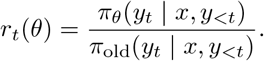

Reinforcement learning was applied only to answer tokens. Image virtual tokens, gene virtual tokens and prompt tokens were used as conditioning context and were not treated as prediction targets. Because this step was intended to improve response style and evidence consistency, the main modality-alignment gains reported in this study are attributed to the three-stage training curriculum rather than to the offline reinforcement learning step.

### 3.7 Implementation details

Training was conducted using the `swift` `pt` framework with a custom registered implementation of SciCore-Omics. Full fine-tuning was used unless otherwise specified. The vision backbone, modality projectors and gene-interface modules were trainable during the corresponding training stages. Mixed-precision training used `bfloat16`.

The main training hyperparameters were as follows: number of epochs = 2, per-device batch size = 1, gradient accumulation steps = 16, learning rate = 5 *×* 10^*−*4^, cosine learning-rate schedule, warmup ratio = 0.1, weight decay = 0.01, maximum sequence length = 2048, evaluation split ratio = 0.05, checkpoint saving every 2,000 steps, logging every 20 steps and random seed = 42. Training was conducted on 8 NVIDIA A100 GPUs with 80 GB memory each.

### 3.8 Transcriptome-to-language evaluation

Transcriptome-to-language generation was evaluated on a held-out validation split of the gene–text paired dataset that was not used for model training. Each model was prompted with transcriptomic input to generate a biological description, which was compared with the corresponding transcriptome-grounded reference description using three complementary metrics. BLEU-4 measures local n-gram precision between generated and reference texts [30]. ROUGE-L evaluates sequence-level structural overlap based on the longest common subsequence [31]. BERTScore F1 computes contextual semantic similarity between generated and reference descriptions in embedding space [32].

Because the reference descriptions were automatically derived from structured labels, marker genes and ssGSEA-derived pathway-enrichment terms, these metrics were interpreted as diagnostics of omics-to-language alignment and structured evidence recovery rather than as standalone evidence of novel biological discovery. Biological grounding was further evaluated through downstream spatial-domain tasks using structured marker-gene and pathway recovery.

### 3.9 Spatial gene expression prediction from histology

To evaluate whether histology-derived representations retained molecular information, we trained a lightweight regression head on top of SciCore-Omics visual embeddings to predict spot-level gene expression from H&E image patches. Experiments were conducted on the normal human heart spatial transcriptomics dataset comprising 39 sections. To avoid information leakage caused by spatial correlations between neighbouring spots within the same tissue section, we used section-level 10-fold cross-validation. Specifically, the 39 sections were partitioned into 10 folds at the section level. In each fold, the model was trained only on spots from the training sections and tested on spots from the held-out section or sections. The prediction targets were the top 50 genes with the highest expression in the training set, and performance was evaluated on the held-out spots from the test sections. Before training and evaluation, all spots were subjected to quality-control filtering, retaining only spots with at least 500 detected genes.

We evaluated two settings. In the frozen-encoder setting, the visual encoder was fixed and only the regression head was trained. In the fine-tuned setting, the regression head and the last six layers of the visual encoder were trainable. Performance was measured using average PCC and MSE across selected highly expressed genes. For each held-out fold, we first computed the PCC for each target gene across all test spots, and then averaged across target genes to obtain the mean PCC for that fold (Extended Data Fig. 3). MSE was computed between the predicted and measured expression matrices. Final results were reported as the average performance across the 10 folds. For all baseline models, we used official implementations whenever possible and retrained them under the same data split, gene panel and preprocessing protocol as our method. To ensure comparability, all models were evaluated on the same held-out spots or sections, and performance was reported using gene-wise PCC and MSE over the same selected genes.

### 3.10 Spatial domain, pathway and marker-gene evaluation

Spatial domain recognition was evaluated on the human dorsolateral prefrontal cortex benchmark. The dataset contained 12 tissue sections, with eight sections used for training or supervised adaptation and four sections held out for evaluation. Manual cortical-layer annotations were used as spatial domain labels.

We evaluated SciCore-Omics in both embedding-based and generative settings. For embedding-based evaluation, frozen model embeddings were extracted from image-only, gene-only or joint image–gene input and passed to a lightweight classification head. Performance was assessed using macro-F1 to account for class imbalance across spatial domains.

For generative evaluation, the model received image-only, gene-only or joint image–gene input and generated a structured biological response. Region labels, pathway labels and marker genes were extracted from the response using deterministic parsing rules based on the required output format. Accuracy was computed separately for region, pathway and marker-gene readouts. The gene readout evaluated whether generated genes appeared in ground-truth marker-gene sets, whereas the pathway readout evaluated consistency between generated pathway labels and ssGSEA-derived pathway annotations. UMAP visualizations were used only for qualitative inspection of embedding organization and were not used for model selection.

### 3.11 Zero-shot tissue-level histopathology classification

We evaluated zero-shot histopathology classification on four public H&E benchmarks: MHIST, BACH, NCT-CRC-HE-100K and PatchCamelyon. For each image, the model was given a dataset-specific closed set of candidate class names and instructed to return exactly one class label. No task-specific fine-tuning, parameter updates or in-context examples were used.

MHIST is a binary colorectal polyp histology dataset containing 3,152 H&E-stained images labelled as hyperplastic polyp or sessile serrated adenoma. BACH is a four-class breast histology dataset containing 400 H&E-stained microscopy images labelled as normal tissue, benign lesion, in situ carcinoma or invasive carcinoma. NCT-CRC-HE-100K is a nine-class colorectal tissue patch dataset containing 100,000 H&E-stained image patches; the independent CRC-VAL-HE-7K validation set was used for evaluation. PatchCamelyon is a binary lymph-node metastasis detection dataset containing 327,680 image patches, with the standard validation split used for evaluation.

For each benchmark, candidate labels were specified using the original dataset class names. Model outputs were post-processed using a deterministic label-matching procedure. Generated responses were converted to lowercase, punctuation was removed and the resulting text was matched against normalized candidate labels and predefined label aliases. When the response contained additional explanatory text, the predicted class was assigned only if exactly one candidate label or alias was present. Responses that could not be mapped unambiguously to a single candidate label were treated as incorrect. Accuracy was computed at the image or patch level as the proportion of correctly classified samples.

For proprietary general-purpose models used as baselines, the same candidate labels and closed-set prompt template were used wherever supported.

### 3.12 Pathology visual question answering

Pathology visual question answering was evaluated on PathVQA, which contains approximately 4,998 pathology images and 32,795 image–question–answer pairs [44]. Evaluation followed the standard separation between open-ended and closed-ended questions. In the fine-tuned setting, the model was trained using image–question–answer triples and evaluated using benchmark-standard answer matching. Closed-ended questions were scored after normalizing yes/no outputs. For open-ended questions, both predictions and reference answers were normalized. Performance was reported as token-level recall, defined as the fraction of reference tokens appearing in the prediction, with multiset matching.

### 3.13 H&E-only case-level pathology workflow

For the pilot whole-slide pathology workflow, only H&E whole-slide images were provided as model input. The cohort included 10 breast cancer cases. Ki-67 immunohistochemistry and pathology reports were withheld from the model and used only as independent references during expert evaluation.

Each slide was first reviewed at low magnification to identify tumour-rich regions and representative tumour–stroma interfaces. High-magnification patches were then selected from these regions for local morphological assessment. The model generated patch-level descriptions covering tumour cellularity, nuclear atypia, chromatin pattern, suspected mitotic activity, necrosis and stromal reaction. Patch-level findings were aggregated into a case-level evidence summary and a proliferation-state interpretation.

Outputs from SciCore-Omics and baseline models were reviewed by pathology experts who were blinded to the source model. The predefined scoring dimensions included accuracy, balanced accuracy, reasoning completeness, evidence quality, molecular consistency and case-level synthesis. Scores were averaged across cases and expert assessments. Because this evaluation included only 10 cases, it was treated as an exploratory proof-of-concept analysis rather than a definitive clinical validation.

### 3.14 Baselines, statistical reporting and reproducibility

#### Baselines

For spatial gene expression prediction, SciCore-Omics was compared with representative histology-based spatial expression prediction methods, including HisToGene, Hist2ST, BLEEP and OmiCLIP [21–24]. For embedding-based spatial domain recognition, comparisons included transcriptomic or spatial representation baselines such as Nicheformer, Geneformer, scGPT, scVI and PCA where applicable. For biomedical and pathology multimodal tasks, comparisons included LLaVA-Med, BioMedGPT, PathGen-LLaVA, Quilt-LLaVA, CellWhisperer and Cell2Sentence where supported by the corresponding input–output format [15–19, 54–57]. For medical visual question answering, additional reported baselines included MMQ, M2I2 and MUMC [58–60].

Baseline models were evaluated using their supported input modalities and task formats. Input– output combinations not supported by a baseline model were not evaluated and were indicated as unsupported in the corresponding figure panels.

#### Statistical reporting and reproducibility

Results are reported on the specified held-out evaluation splits for each task. For histopathology benchmark comparisons, mean and standard deviation summarize variation across the four benchmark datasets rather than repeated model-training runs. For spatial transcriptomics experiments, spot-level predictions were aggregated within held-out sections, and section-level splits were used to reduce spatial leakage. For experiments where repeated full-model training, bootstrap resampling or seed-based uncertainty estimation was not performed, confidence intervals are not reported.

To support reproducibility, dataset sources, preprocessing procedures, split strategies, model architecture and principal training hyperparameters are described above. Public datasets and download locations are listed in the Data availability section and Supplementary Table 1. The codebase and training configuration are provided in the Code availability section. Source data for plotted aggregate results are provided where permitted by dataset licenses and institutional data-governance restrictions.

### 3.15 Model parameter reporting

To contextualize the computational scale of SciCore-Omics and the evaluated baselines, we reported model parameter counts whenever they were publicly available or could be derived from released model configurations. SciCore-Omics contains approximately 8.5 billion parameters, including the language-centred multimodal backbone, histology visual branch, Nicheformer-based gene encoder, Q-Former module and modality projection layers. For open-source baseline models, parameter counts were obtained from the corresponding papers, model cards or official implementations. For proprietary models, including GPT-4o and GPT-5, parameter counts are not publicly disclosed and are therefore marked as not available. For classical or configuration-dependent methods, including PCA and scVI, model size was reported as not applicable or implementation-dependent. The parameter counts of SciCore-Omics and all evaluated baselines are summarized in Supplementary Table 2.

## 4 Data availability

All public datasets and resources used in this study are publicly available and can be accessed from: STimage-1K4M [61] (https://github.com/JiawenChenn/STimage-1K4M; https://huggingface.co/datasets/jiawennnn/STimage-1K4M), PubMed (https://pubmed.ncbi.nlm.nih.gov/), PubMed Central (https://pmc.ncbi.nlm.nih.gov/), GEO (https://www.ncbi.nlm.nih.gov/geo/), 10x Genomics public datasets (https://www.10xgenomics.com/datasets), PathGen-1.6M (https://huggingface.co/datasets/jamessyx/PathGen; https://github.com/PathFoundation/PathGen-1.6M), UltraChat (https://huggingface.co/datasets/openbmb/UltraChat), Flickr30k (https://opendatalab.com/OpenDataLab/Flickr30k/download), human DLPFC spatial transcriptomics data through spatialLIBD and spatialDLPFC (https://research.libd.org/spatialLIBD/; https://github.com/LieberInstitute/spatialDLPFC), the human heart spatial transcriptomics dataset (https://www.heartcellatlas.org/), MHIST (https://bmirds.github.io/MHIST/), BACH (https://iciar2018-challenge.grand-challenge.org/Dataset/), NCT-CRC-HE-100K and CRC-VAL-HE-7K (https://zenodo.org/records/1214456), PatchCamelyon (https://patchcamelyon.grand-challenge.org/; https://www.tensorflow.org/datasets/catalog/patch_camelyon) and PathVQA (https://github.com/KaveeshaSilva/PathVQA). We summarize the dataset statistics and download information in Supplementary Table 1. The internal breast cancer H&E and immunohistochemistry whole-slide image cohort used for the pilot case-level workflow is not publicly released because of patient privacy and institutional data-governance restrictions. De-identified aggregate results and expert-evaluation scores used to generate the corresponding figure panels are provided as source data where permitted.

## 5 Code availability

The pretrained model, as well as source code for training, inference and data preprocessing, can be accessed at https://github.com/OpenBMB/Scicore-Omics. A web-based demonstration platform for SciCore-Omics is available at https://huggingface.co/spaces/Alkaidxxy/SciCore-Omics.

## 6 Acknowledgements

We thank our collaborators and laboratory members for helpful discussions and technical support. We also acknowledge the developers and maintainers of MiniCPM-V, Nicheformer and the public datasets and benchmark resources used in this study.

## 8 Author contributions

X.X. and Z.Z. conceptualized the study and designed the overall framework. X.X. and Y.F.L. developed the methodology, implemented the software, performed data processing, conducted the experiments and interpreted the results. Z.B.Y., Y.J.X. and J.M.Y. contributed data resources and provided clinical and pathological expertise. Z.Z., Y.K.Y., Z.Y.L., Z.H.L., J.M.Y., Y.L., L.H.X. and F.C.H. supervised the work and provided scientific guidance. X.X. wrote the original draft. All authors reviewed and edited the manuscript, provided critical feedback, and approved the final manuscript.

## 9 Ethics declarations

### Ethics approval and consent to participate

The use of the internal breast cancer cohort was approved by the Ethics Committee of National Cancer Center/Cancer Hospital, Chinese Academy of Medical Sciences and Peking Union Medical College (approval no. 25/077-5023). The study was conducted using de-identified retrospective data in accordance with relevant institutional guidelines and regulations. The requirement for written informed consent was waived by the ethics committee.

## Extended Data Figures

**Extended Data Fig. 1:**
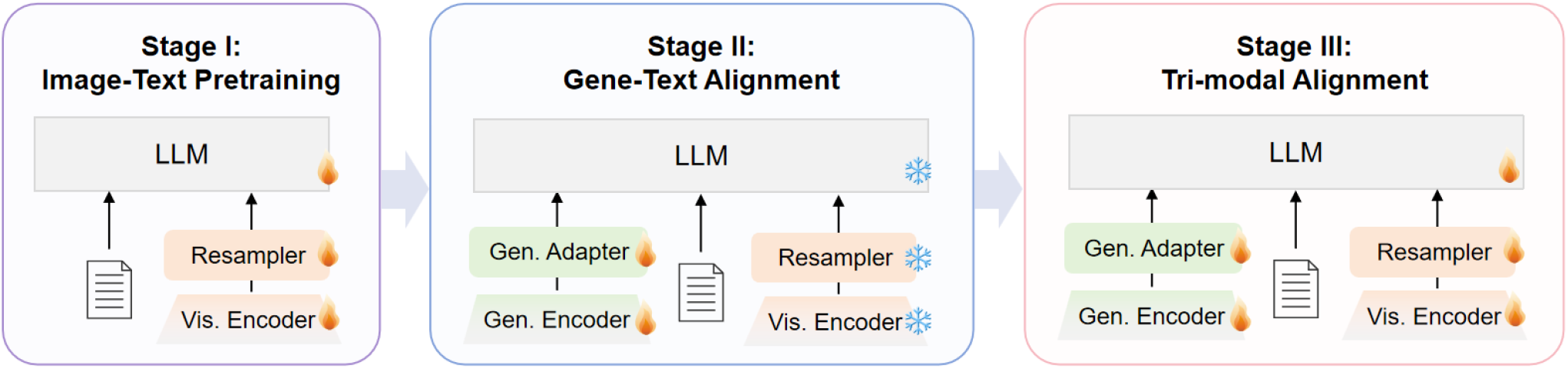
Module-level implementation of the progressive multimodal alignment strategy. Stage I performs pathology-oriented image–text pretraining by optimizing the vision encoder, resampler and language model components. Stage II introduces the gene encoder and gene adapter to align transcriptomic representations with the language token space. Gen. Adapt denotes the gene adapter, implemented as a Gene Q-Former followed by a projector. Stage III jointly optimizes image, gene and text representations using paired tri-modal data. Flame icons indicate trainable modules, whereas snowflake icons indicate frozen modules.

**Extended Data Fig. 2:**
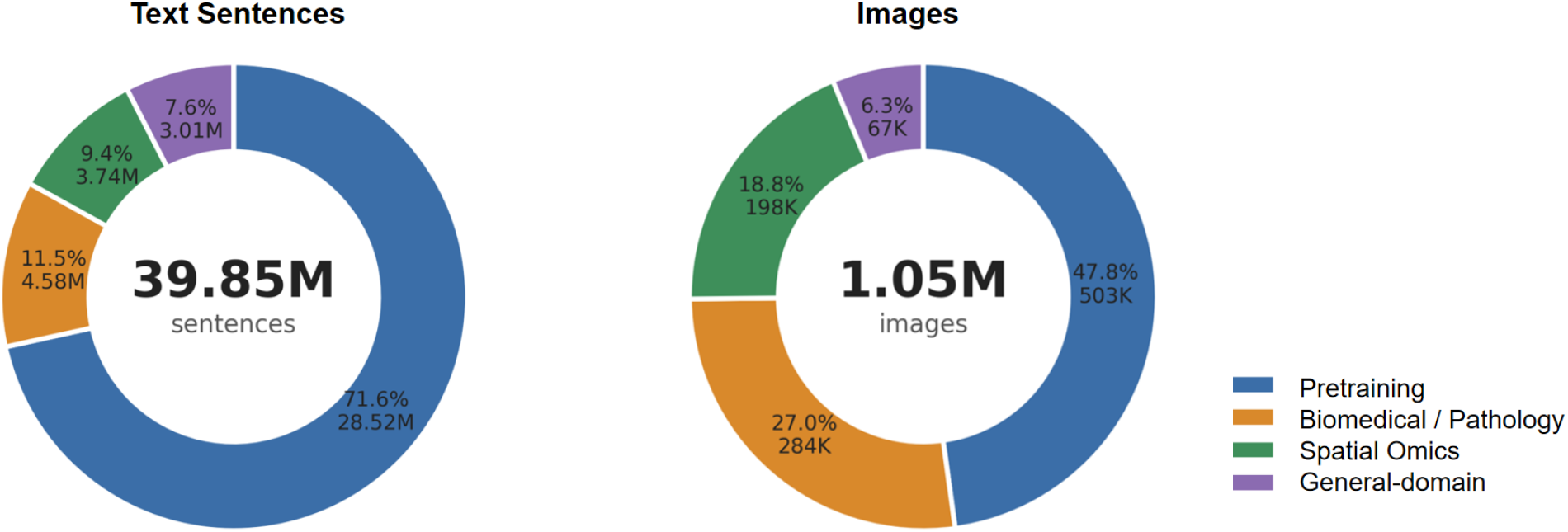
Composition of the SciCore-Omics training corpus. Summary of text-only, image–text, gene–text and image–gene–text data used for progressive multimodal alignment. The corpus included approximately 1.05 million images, 39.85 million text sentences and 151,182 tri-modal spatial transcriptomics spots across multiple tissue types.

**Extended Data Fig. 3:**
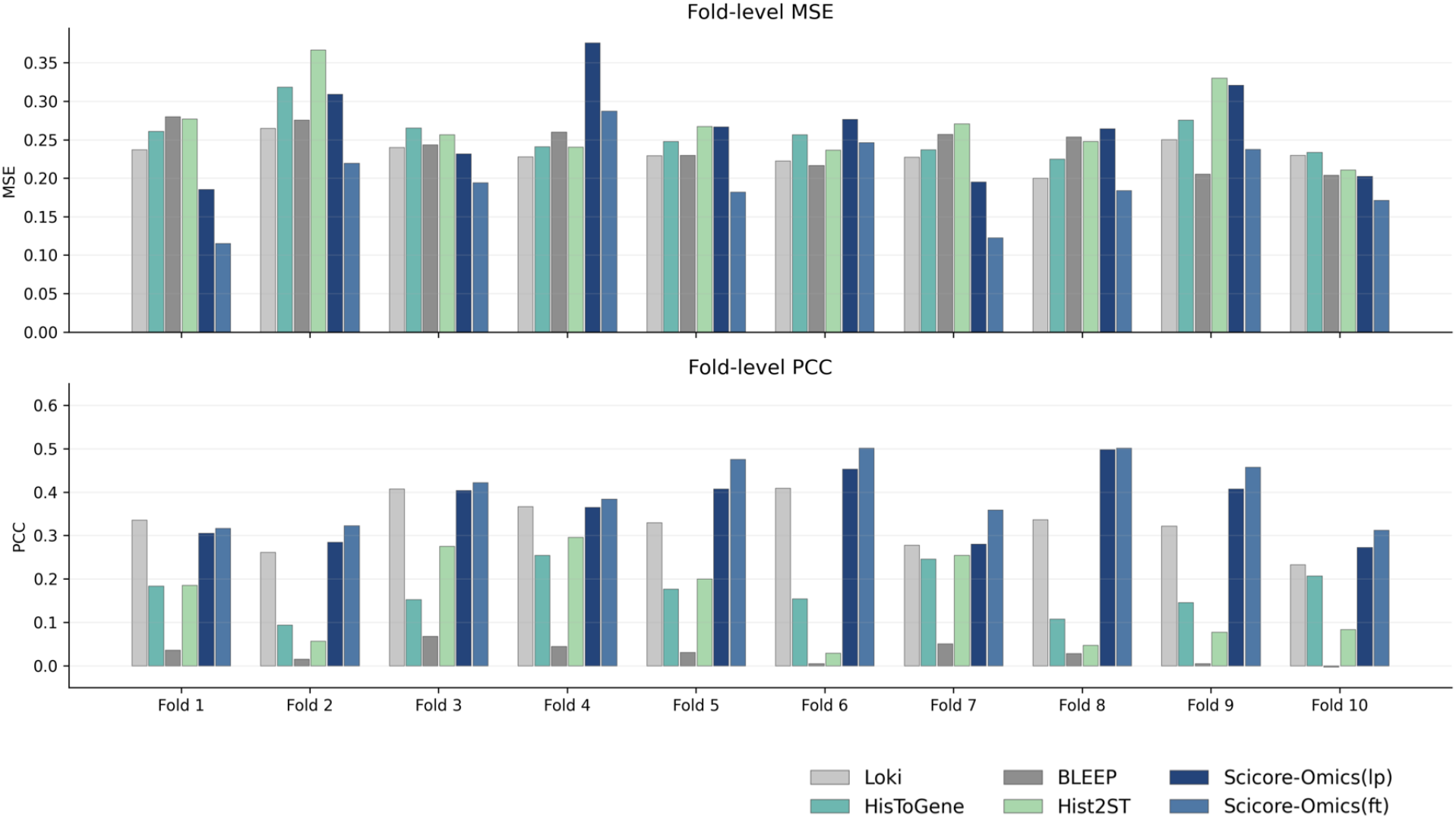
Cross-validation results for histology-based gene expression prediction. Supplementary performance of SciCore-Omics on the normal human heart spatial transcriptomics dataset using section-level 10-fold cross-validation. PCC and MSE were computed for the top 50 highly expressed genes on held-out sections under frozen-encoder and fine-tuned settings.

**Extended Data Fig. 4:**
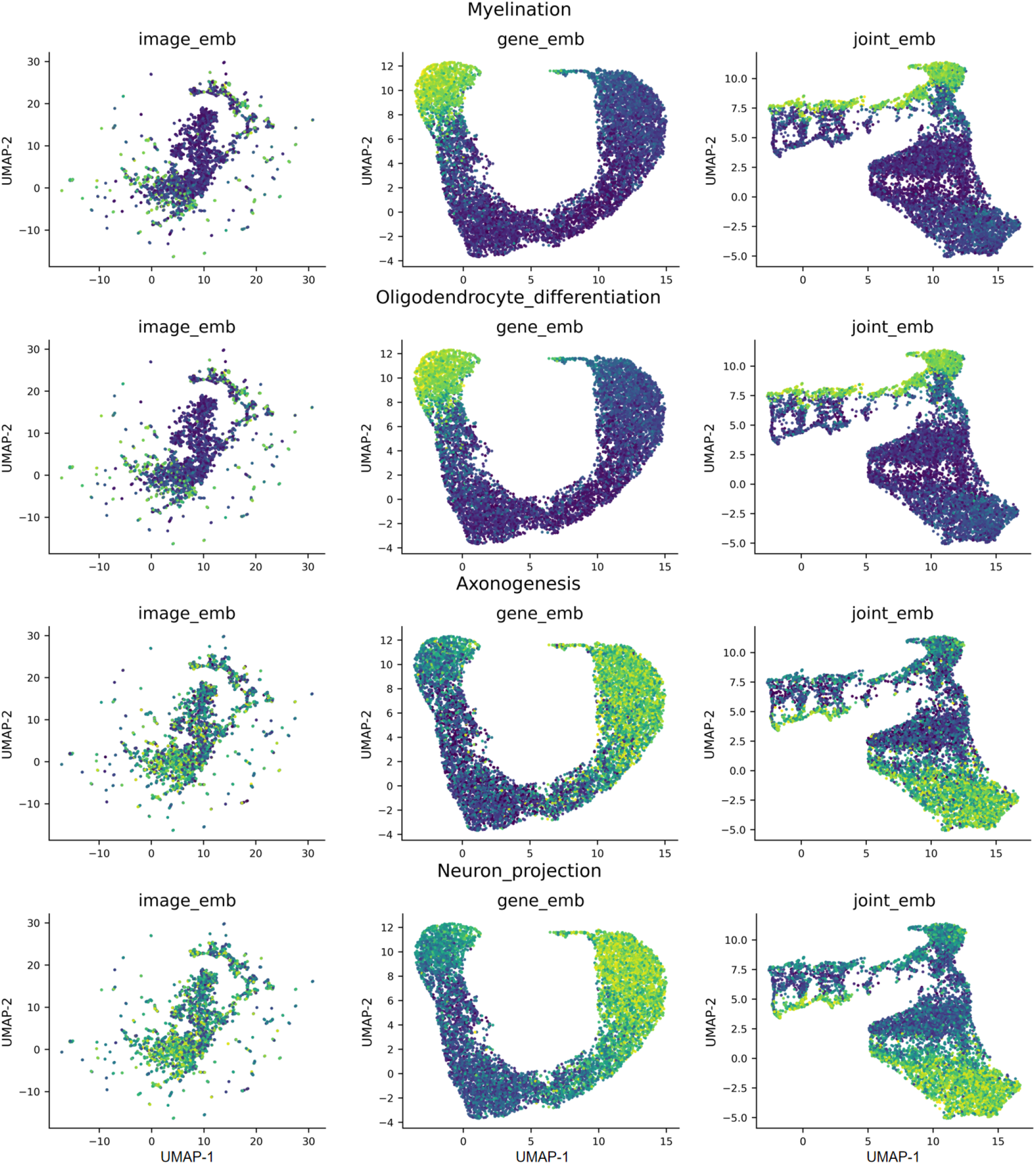
UMAP visualization of pathway activity across SciCore-Omics embeddings. UMAP projections of image-only, gene-only and joint image–gene embeddings from the DLPFC spatial transcriptomics dataset, coloured by pathway activity scores for representative pathways, including *Myelination, Oligodendrocyte differentiation, Axonogenesis* and *Neuron projection*. Joint image–gene embeddings showed clearer pathway-associated organization than single-modality embeddings.

**Extended Data Fig. 5:**
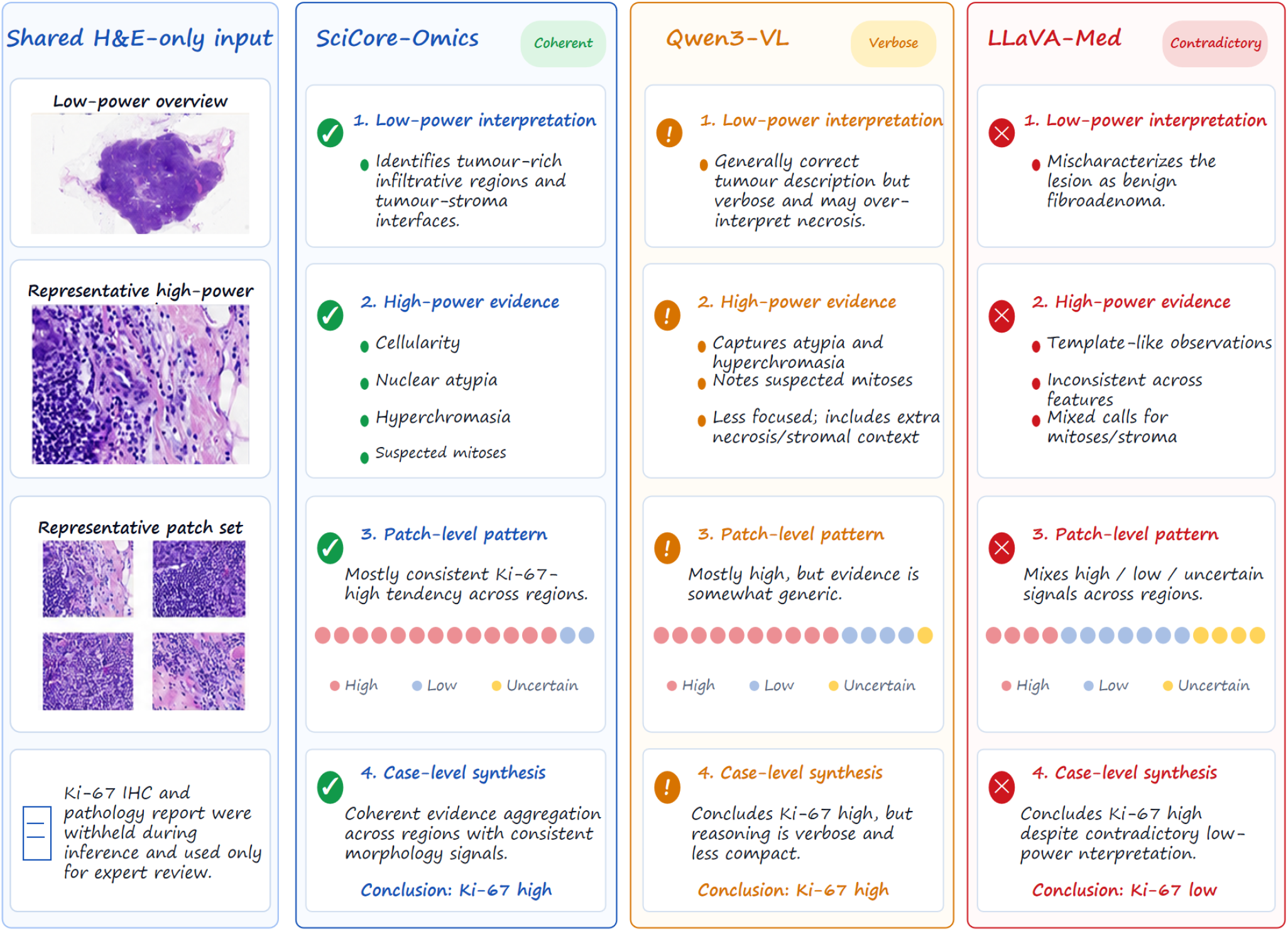
Representative H&E-only case-level reasoning comparison. All models received the same H&E-only breast cancer case, while Ki-67 immunohistochemistry and pathology reports were withheld during inference. SciCore-Omics produced coherent evidence supporting a Ki-67-high conclusion; Qwen3-VL was relevant but verbose, whereas LLaVA-Med showed inconsistent reasoning and concluded Ki-67 low status. Coloured dots indicate patch-level high, low or uncertain Ki-67 tendencies.

