## Supplementary Information for "SciCore-Omics: a tri-modal foundation model unifying histology, spatial transcriptomics and language for spatial biology"

### Supplementary Tables

Table S1: **Supplementary Table 1 – Dataset statistics and download information.** Public datasets and auxiliary resources used for model development, training or evaluation are listed together with their modality, role in this study, associated task or figure, dataset statistics and access information. Internal clinical whole-slide images were used only for the pilot case-level workflow and are not publicly released because of patient privacy and institutional data-governance restrictions.

| **No.** | **Dataset or resource** | **Modality** | **Role in this study** | **Task or figure** | **Dataset statistics** | **Download, accession or availability** |
|---|---|---|---|---|---|---|
| 1 | STimage-1K4M | H&E histology, spatial gene expression and spatial coordinates | Core tri-modal spatial transcriptomics resource used to construct spot-level image–gene–text triplets and to support transcriptome-to-language evaluation | Stage III training; Fig. 2 | 1,149 spatial transcriptomics slides and more than 4 million spatial spots with paired histology images, gene-expression profiles and spatial coordinates | `https://github.com/JiawenChenn/STimage-1K4M`; `https://huggingface.co/datasets/jiawennnn/STimage-1K4M`. Approximately 150,000 spot-level triplets were used after filtering in this study. |
| 2 | PubMed and PubMed Central pathology image–text resources | Pathology figures, captions, article text and metadata | Pathology semantic grounding for image–language alignment | Stage I training | Approximately 250,000 pathology image–text pairs curated in this study from publicly available biomedical literature | PubMed: `https://pubmed.ncbi.nlm.nih.gov/`; PubMed Central: `https://pmc.ncbi.nlm.nih.gov/`. Article identifiers, figure identifiers and filtering metadata are documented in the project records and are released where licensing permits. |
| 3 | GEO transcriptomic resources | Bulk, single-cell or spatial transcriptomic profiles with sample metadata | Gene–text alignment using transcriptome-derived pathway, gene and tissue descriptions | Stage II training | Public transcriptomic studies curated from GEO; used together with 10x Genomics resources to construct approximately 50,000 gene–text pairs | `https://www.ncbi.nlm.nih.gov/geo/`. Public GEO records were accessed through NCBI GEO. Curated accession metadata and processing scripts are provided in the associated project repository or from the corresponding authors where redistribution is permitted. |
| 4 | 10x Genomics public datasets | Single-cell and spatial gene-expression data | Auxiliary public transcriptomic resources for gene–text alignment | Stage II training | Public 10x Genomics datasets spanning multiple tissues, assays and sample types; used as part of the approximately 50,000 gene–text pair collection | `https://www.10xgenomics.com/datasets`. Dataset names, assay versions and filtering scripts are documented in the project records and are made available through the associated repository where permitted by dataset terms. |
| 5 | PathGen-1.6M | Pathology image–caption pairs | Auxiliary pathology image–text training to preserve and improve pathology vision–language capability | Stage III and Stage IV auxiliary training | 1.6 million generated pathology image–caption pairs derived from representative whole-slide image patches | `https://huggingface.co/datasets/jamessyx/PathGen`; `https://github.com/PathFoundation/PathGen-1.6M`. A sampled subset was used in this study subject to dataset licence and source-slide redistribution restrictions. |

Continued on next page

Table S1: **Supplementary Table 1** continued.

| No. | Dataset or resource | Modality | Role in this study | Task or figure | Dataset statistics | Download, accession or availability |
|---|---|---|---|---|---|---|
| 6 | UltraChat | Text-only multi-turn instruction data | Auxiliary instruction-following data for preserving general language capability during multimodal training | Stage III and Stage IV auxiliary training | Large-scale open-source multi-round dialogue dataset | `https://huggingface.co/datasets/openbmb/UltraChat`. A sampled subset was used as general instruction data. |
| 7 | Flickr30k | Natural images and captions | Auxiliary image–text data used to preserve general visual–language capability | Stage III auxiliary training | 31,783 images with 158,915 English captions | `https://opendatalab.com/OpenDataLab/Flickr30k/download`; additional entity annotations: `https://github.com/bryanplummer/flickr30k_entities`. |
| 8 | Human DLPFC spatial transcriptomics dataset | H&E histology, 10x Genomics Visium spatial transcriptomics and cortical-layer annotations | Spatial-domain, pathway and marker-gene evaluation | Fig. 4 | Human dorsolateral prefrontal cortex spatial transcriptomics dataset generated from postmortem human brain samples; spatialLIBD provides processed spot-level data and layer annotations | spatialLIBD: `https://research.libd.org/spatialLIBD/`; spatialDLPFC repository: `https://github.com/LieberInstitute/spatialDLPFC`. Dataset sections were used according to the spatial-domain benchmark protocol. |
| 9 | Human Heart Cell Atlas spatial transcriptomics dataset, E-MTAB-12975 | H&E histology and Visium spatial transcriptomics | Spatial gene-expression prediction from histology | Fig. 3 | Human heart spatial transcriptomics generated as part of the spatially resolved multiomics study of human cardiac niches | BioStudies/ArrayExpress accession E-MTAB-12975: `https://www.ebi.ac.uk/biostudies/arrayexpress/studies/E-MTAB-12975`. Normal human heart sections were used for image-to-gene prediction. |
| 10 | MHIST | H&E histopathology image patches | Zero-shot histopathology classification | Fig. 5 | 3,152 H&E-stained fixed-size images of colorectal polyps, each 224 × 224 pixels; binary labels correspond to hyperplastic polyp and sessile serrated adenoma | `https://bmirds.github.io/MHIST/`. The public benchmark images were used for closed-set zero-shot classification. |
| 11 | BACH | Breast H&E microscopy images and whole-slide images | Zero-shot breast histopathology classification and confusion-matrix analysis | Fig. 5 | 400 microscopy images distributed equally across normal, benign, in situ carcinoma and invasive carcinoma classes; the full challenge dataset also includes whole-slide images | `https://iciar2018-challenge.grand-challenge.org/Dataset/`. The microscopy image classification subset was used in this study. |
| 12 | NCT-CRC-HE-100K and CRC-VAL-HE-7K | Colorectal H&E image patches | Zero-shot colorectal histopathology classification | Fig. 5 | NCT-CRC-HE-100K contains 100,000 H&E image patches. CRC-VAL-HE-7K contains 7,180 image patches from 50 patients without patient overlap with NCT-CRC-HE-100K. Images are 224 × 224 pixels at 0.5 microns per pixel | Zenodo record: `https://zenodo.org/records/1214456`. The benchmark was used for nine-class colorectal tissue classification. |

Continued on next page

Table S1: **Supplementary Table 1** continued.

| No. | Dataset or resource | Modality | Role in this study | Task or figure | Dataset statistics | Download, accession or availability |
|---|---|---|---|---|---|---|
| 13 | PatchCamelyon | Lymph-node H&E image patches | Zero-shot metastatic tissue classification | Fig. 5 | 327,680 colour images, each 96 × 96 pixels, extracted from histopathological scans of lymph-node sections and annotated with binary metastatic-tissue labels | Grand Challenge: `https://patchcamelyon.grand-challenge.org/`; TensorFlow Datasets: `https://www.tensorflow.org/datasets/catalog/patch_camelyon`. Official splits were used for evaluation. |
| 14 | PathVQA | Pathology images and visual question–answer pairs | Pathology visual question answering | Fig. 6a | 4,998 pathology images and 32,799 question–answer pairs generated from pathology educational resources | `https://github.com/KaveeshaSilva/PathVQA`. Official train, validation and test splits were used where available. |
| 15 | Internal breast cancer H&E and immunohistochemistry whole-slide image cohort | H&E WSI, Ki-67 IHC, PR IHC, HER2 IHC and pathology reports | Pilot H&E-only case-level pathology reasoning and expert assessment | Fig. 6b–d | Ten de-identified retrospective breast cancer cases were used for the pilot case-level workflow. H&E whole-slide images were provided as model input; Ki-67 IHC, PR IHC, HER2 IHC and pathology reports were used only as independent references for expert assessment. | Not publicly released because of patient privacy and institutional data-governance restrictions. The retrospective cohort was approved by the Ethics Committee of National Cancer Center/Cancer Hospital, Chinese Academy of Medical Sciences and Peking Union Medical College, approval no. 25/077-5023. De-identified aggregate results and expert-evaluation scores are provided with the paper where permitted. |

Table S2: **Supplementary Table 2 – Model parameter counts for SciCore-Omics and evaluated baselines.** Parameter counts are reported when publicly available from the corresponding paper, model card, official repository or released configuration. For proprietary models, classical methods or methods whose size depends on a specific implementation, the parameter count is marked as not disclosed, not applicable or implementation-dependent. Counts refer to total model scale unless otherwise stated.

| No. | Model or method | Model category | Used in this study | Reported parameter count | Reporting basis and notes |
|---|---|---|---|---|---|
| 1 | SciCore-Omics | Tri-modal histology–omics–language foundation model | Main model | Approximately 8.5 billion | Estimated from the implementation used in this study. The count includes the language-centred multimodal backbone, histology visual branch, Nicheformer-based gene encoder, Gene Q-Former module and modality projection layers. |
| 2 | HisToGene | Histology-to-gene-expression prediction model | Fig. 3 gene-expression prediction baseline | Implementation-dependent; not reported as a single public total | The original method is a task-specific neural model for predicting spatial gene expression from histology. The total number of parameters depends on the selected image backbone, transformer or regression-head configuration and target gene panel. |
| 3 | Hist2ST | Histology-to-spatial-transcriptomics prediction model | Fig. 3 gene-expression prediction baseline | Implementation-dependent; not reported as a single public total | The model combines image features, spatial information and graph or transformer components. The effective model size varies with the released configuration, graph construction and output gene set used for a given benchmark. |
| 4 | BLEEP | Bi-modal image–expression contrastive model | Fig. 3 gene-expression prediction baseline | Implementation-dependent; not reported as a single public total | The total count depends on the image encoder, expression encoder and projection heads used in the reproduced configuration. Therefore, we report it as implementation-dependent rather than assigning a single architecture-agnostic value. |
| 5 | OmiCLIP | Visual–omics foundation model | Fig. 3 gene-expression prediction baseline | Not publicly disclosed as a single total in the source used for comparison | OmiCLIP is a CLIP-style visual–omics model trained on paired tissue images and transcriptomic data. Because the total checkpoint-level parameter count is not consistently reported, it is listed as not publicly disclosed. |
| 6 | Nicheformer | Single-cell and spatial-omics foundation model | Fig. 4 embedding baseline and SciCore-Omics gene-encoder component | 49.3 million | Reported for the published Nicheformer architecture with a 1,500-token context length, 12 transformer encoder units, 16 attention heads per layer and a 512-dimensional embedding. |
| 7 | Geneformer | Single-cell transcriptomic foundation model | Fig. 4 embedding baseline | 10 million for Geneformer-V1; 104 million or 316 million for Geneformer-V2 variants | The public Geneformer repository and model cards provide multiple checkpoints. The exact parameter count should be matched to the specific checkpoint used in an evaluation run. |
| 8 | scGPT | Single-cell multi-omics generative foundation model | Fig. 4 embedding baseline | Approximately 53 million for the commonly used pretrained checkpoint | The parameter count corresponds to the released pretrained scGPT model commonly used for single-cell and multi-omics representation learning. |

Continued on next page

Table S2: **Supplementary Table 2** continued.

| No. | Model or method | Model category | Used in this study | Reported parameter count | Reporting basis and notes |
|---|---|---|---|---|---|
| 9 | scVI | Deep generative model for single-cell transcriptomics | Fig. 4 embedding baseline | Implementation-dependent | scVI is a configurable variational autoencoder rather than a fixed-size foundation model. Its parameter count depends on the input gene dimension, latent dimension, hidden width and number of layers. |
| 10 | PCA | Classical dimensionality-reduction method | Fig. 4 embedding baseline | Not applicable | PCA does not have neural-network parameters. The number of fitted loadings depends on the number of input features and retained principal components. |
| 11 | LLaVA-Med | Biomedical vision–language model | Fig. 5 zero-shot classification baseline and Fig. 6 case-level reasoning baseline | 7B or 13B language-backbone variants, plus vision encoder and projector | LLaVA-Med is built by biomedical instruction tuning of LLaVA-style models. The original work reports 7B and 13B variants; the total multimodal count additionally includes the visual encoder and projection layers. |
| 12 | BioMedGPT | Unified biomedical generative model | Fig. 6a PathVQA baseline | 33M, 93M and 182M variants; BioMedGPT-B has 182M parameters | The BioMedGPT paper explicitly reports small, medium and base variants. The base model, BioMedGPT-B, is the 182M-parameter version. |
| 13 | PathGen-LLaVA | Pathology-specific large multimodal model | Fig. 5 zero-shot classification baseline | Approximately 13B language backbone, plus PathGen-CLIP-L vision encoder and projector | PathGen-LLaVA follows the LLaVA-v1.5-13B framework and replaces the vision encoder with PathGen-CLIP-L. The total multimodal size is therefore larger than the 13B language backbone alone. |
| 14 | Quilt-LLaVA | Histopathology-specific vision–language model | Fig. 5 zero-shot classification baseline | Approximately 13B language backbone, plus vision encoder and projector | Quilt-LLaVA is trained in the LLaVA family using histopathology instruction data. Because the exact merged checkpoint size depends on the released configuration, we report the language-backbone scale and note the additional multimodal components. |
| 15 | CellWhisperer | Transcriptome–language and chat-based single-cell model | Biomedical multimodal baseline where supported | Approximately 7.3B for the Mistral-7B chat backbone, plus transcriptome–text embedding modules | The CellWhisperer chat model builds on Mistral 7B. The contrastive transcriptome–text embedding model can also be used independently, so the exact total depends on whether the chat model, embedding model or combined system is evaluated. |
| 16 | Cell2Sentence | LLM-based single-cell transcriptome language model | Omics–language baseline where supported | 410M in commonly released C2S-Pythia examples; C2S-Scale family ranges from 410M to 27B | Cell2Sentence adapts language models to transcriptomic "cell sentences". Parameter count is checkpoint-specific; recent C2S-Scale models include 410M, 1B, 2B, 9B and 27B variants. |
| 17 | GPT-4o | Proprietary general-purpose multimodal model | Fig. 5 zero-shot classification baseline | Not publicly disclosed | OpenAI has not publicly released a verified parameter count for GPT-4o. We therefore mark the parameter count as unavailable rather than relying on third-party estimates. |
| 18 | GPT-5 | Proprietary general-purpose multimodal and reasoning model | Fig. 5 zero-shot classification baseline | Not publicly disclosed | OpenAI has not publicly released a verified parameter count for GPT-5. The model is therefore listed as not publicly disclosed. |

Continued on next page

Table S2: **Supplementary Table 2** continued.

| No. | Model or method | Model category | Used in this study | Reported parameter count | Reporting basis and notes |
|---|---|---|---|---|---|
| 19 | Qwen3-VL | Open-weight multimodal large language model family | Fig. 6 case-level reasoning baseline | Dense variants include 2B, 4B, 8B and 32B; MoE variants include 30B-A3B and 235B-A22B | Qwen3-VL is released as a family of models. The precise parameter count should be reported according to the evaluated checkpoint; A3B and A22B denote approximate activated parameters for the MoE variants. |
| 20 | MMQ | Medical visual question answering model | Fig. 6a PathVQA baseline | Implementation-dependent; not reported as a single public total | MMQ is a pre-large-language-model Med-VQA architecture based on multiple meta-model quantifying. Its size depends on the selected visual extractor, language encoder and classifier configuration. |
| 21 | M2I2 | Self-supervised medical vision–language pretraining model | Fig. 6a PathVQA baseline | Implementation-dependent; not reported as a single public total | M2I2 combines masked image modeling and contrastive learning for MedVQA. The effective parameter count depends on the selected visual and textual encoders and downstream head. |
| 22 | MUMC | Medical vision–language pretraining model with unimodal and multimodal contrastive losses | Fig. 6a PathVQA baseline | Implementation-dependent; not reported as a single public total | MUMC extends medical vision–language pretraining with unimodal and multimodal contrastive objectives. The total count is configuration-dependent and is not consistently reported as a single public value. |
